# Structure and function of a fungal AB toxin-like chimerolectin involved in anti-nematode defense

**DOI:** 10.1101/2025.08.11.669643

**Authors:** Stefanie S. Schmieder, Gabriele Cordara, Flore Kersten, Kevin Steiner, Clara H. Samin, David F. Plaza, Ahmad A. Ahmad, Andreas Boeggild, Jesper L. Karlsen, Blanka O. Sokolowska, Thomas Boesen, Ute Krengel, Markus Künzler

## Abstract

Fungal defense against predators largely relies on protein toxins, many of which are lectins. We previously showed that the production of the nematotoxin CCTX2 is upregulated in the Agaricomycete *Coprinopsis cinerea* upon predation by nematodes. Here, we classify CCTX2 as the founding member of a family of fungal chimerolectins. Cryo-EM analysis to 3.2 Å resolution reveals five domains. The four N-terminal β-trefoil fold (BTF) domains cradle a C-terminal domain, which exhibits a novel α+β protein fold. Mutational analysis shows that both N-terminal and C-terminal domains are required for nematotoxicity. While the biochemical function of the C-terminal domain remains unclear, the first two BTF domains enable CCTX2 to bind to glycosphingolipids with LacNAc or LacdiNAc glycoepitopes on nematode intestinal epithelial cells. Experiments in the model nematode *Caenorhabditis elegans* demonstrate that the chimerolectin CCTX2 exploits the endocytic and retrograde trafficking machinery of the target cell to exert its toxicity and obtain access to the yet-to-be-identified intracellular target of the non-lectin domain. The structure and mode of action of CCTX2 is reminiscent of bacterial and plant AB toxins.

## INTRODUCTION

Proteinaceous toxins are involved in many inter-organismal conflicts and are especially prevalent in plants and bacteria (Aravind et al, 2012; Zhang et al, 2012). AB toxins are one form of such defense toxins and are formed by two protein components: the A subunit (**A**ctive subunit), responsible for exerting toxicity inside the cell, and B subunits (**B**inding subunits), responsible for cell binding and entry (Song, 2022). Ricin (plants), Shiga and cholera toxin (bacteria) are considered prime examples of this class of toxins (Zhang et al, 2012; Melton-Celsa, 2014; Clemens et al, 2017; Polito et al, 2019). AB toxins are found in nature with varying subunit stoichiometries, ranging from AB (*e.g.,* ricin), to AB_2_ (*e.g.,* cytolethal distending toxin, CDT), AB_5_ (*e.g.,* Shiga and cholera toxins) or A_2_B_5_ (*e.g.,* typhoid toxin) (Pickett & Whitehouse, 1999; Vanden Broeck et al, 2007; Song et al, 2013a; Melton-Celsa, 2014; Clemens et al, 2017; Polito et al, 2019).

The B subunit often recognizes a glycoepitope with high specificity, thus classifying it as a sugar-binding protein, or lectin. This subunit mediates the binding of the AB-toxin to a glycoconjugate on the target cell surface and triggers the entry of the toxin into the target cell by endocytosis (Mancheño et al, 2005; Manna et al, 2017). Once inside, the A subunit exerts its toxicity by binding to and/or modifying one or more intracellular target molecules (Teter, 2013). Catalytic activities of AB toxins include ADP-ribosyltransferase (ART, cholera toxin) and rRNA *N*-glycosidase activity (ricin and Shiga toxin) (Sánchez & Holmgren, 2011; Melton-Celsa, 2014; Polito et al, 2019).

The sessile lifestyle and the high concentration of macronutrients render fungi an attractive food source for the soil-grazing fauna. To protect themselves from being foraged by *e.g.* nematodes, fungi employ a combination of secondary metabolites and a protein-based defense system; the latter including several lectins (Singh et al, 2010; Bleuler-Martínez et al, 2011; Kempken, 2011; Varrot et al, 2013). Fungivorous nematodes pierce fungal cells, and aspirate their cytoplasmic content; consistently, defense lectins are cytoplasmically localized (Künzler, 2015). After ingestion, the lectin travels through the digestive tract of nematodes to eventually bind to a cell surface receptor and exert its toxic effect (Künzler, 2015).

Lectins include mero-(monomeric) and hololectins (multimeric), exclusively formed by sugar-binding domains, and chimerolectins, where lectin and non-lectin domains coexist or assemble in the same protein (Peumans & Van Damme, 1995; Peumans et al, 2001). Some chimerolectins are functionally analogous to AB toxins. Studies of the fungal chimerolectin *Marasmius oreades* agglutinin (MOA) suggest that the lectin domain acts as a B subunit and the non-lectin domain as an A subunit (Cordara et al, 2011; Wohlschlager et al, 2011; Cordara et al, 2016). The toxicity mechanism of MOA includes binding to glycosphingolipids exposed on the surface of the target cell as well as endocytosis, retrograde trafficking, and (putative) modification of an intracellular target (Cordara et al, 2011; Wohlschlager et al, 2011; Cordara et al, 2014; Künzler, 2015; Juillot et al, 2016). MOA exerts strong toxicity against nematodes, including the bacterivorous model organism *Caenorhabditis elegans*, and is thought to be part of the chemical defense of the host *M. oreades* against these micropredators (Wohlschlager et al, 2011; Tayyrov et al, 2018).

CCTX2 was previously identified as a nematotoxic defense protein in the Agaricomycete *Coprinopsis cinerea*, based on the induced expression of the *cctx2* gene in the vegetative mycelium of the fungus in the presence of fungivorous nematodes (Plaza et al, 2016). In the present article, we performed a structure/function analysis of the CCTX2 protein toxin. Within the *C. cinerea* genome, we additionally identified the genes of two CCTX2 paralogs, which we termed *cctx1* and *cctx3*. The three genes differ in their expression pattern regarding sexual development and bacterial or nematode challenge; however, we demonstrate that the encoded proteins all exert strong nematotoxicity and show the same domain partition: four N-terminal β-trefoil fold (BTF) domains followed by a C-terminal domain of unknown function (DUF). The structure of CCTX2, determined to 3.2 Å resolution by single-particle cryo–electron microscopy (cryo-EM), revealed that the BTF domains form a cradle carrying the C-terminal domain. The C-terminal domain adopts a novel α+β protein fold, not represented in any protein structures deposited in the Protein Data Bank (PDB). Deletion of the first two BTF domains of CCTX2 leads to the loss of binding of the chimerolectin to LacNAc and LacdiNAc glycoepitopes. This, together with the presence of experimental density at the carbohydrate-binding sites in the two N-terminal BTF domains, suggests that the nematotoxicity of CCTX2 depends on the binding of the protein to LacdiNAc-containing glycosphingolipids (GSLs) on the surface of nematode intestinal epithelial cells. Upon binding, CCTX2 undergoes endocytosis and retrograde trafficking of the protein-GSL complexes in these cells. The nematotoxicity of CCTX2 also depends on the presence and integrity of the C-terminal, non-lectin domain. The intracellular target and the biochemical function of this domain remain unknown. Based on orthologues encoded in the genome of other Agaricomycetes and some ascomycetes, we conclude that CCTX2 is the founding member of a family of fungal defense effectors against fungivorous nematodes.

## RESULTS

### CCTX2 belongs to a family of fungal nematotoxins

A query on the *Coprinopsis cinerea* genome (JGI Mycocosm *Coprinopsis cinerea* AmutBmut v2) using the *cctx2* gene sequence (gene model ID CopciAB_369589.T0) revealed the existence of two paralogues, aptly designated as *cctx1* and *cctx3* (gene model IDs CopciAB_369594.T0 and CopciAB_369576.T0, respectively), which are located on the same chromosome in close neighborhood of the *cctx2* gene (Fig. 1A). According to published data (Muraguchi et al, 2015; Tayyrov et al, 2018; Kombrink et al, 2019) (Fig. S1A-B), the three paralogues differ significantly in their expression patterns: *cctx2* is expressed at low levels in the vegetative mycelium and is induced upon predation by the fungivorous nematode *Aphelenchus avenae* but not upon bacterial challenge. Expression of *cctx1* is low also in vegetative mycelium, but the gene is strongly upregulated upon nematode challenge and in stage 1 primordia during sexual development, whereas *cctx3* is strongly expressed in the vegetative mycelium and downregulated during sexual development as well as upon nematode and bacterial challenge. A BLAST search using the full-length CCTX1, CCTX2 and CCTX3 sequences (JGI Mycocosm *Coprinopsis cinerea* AmutBmut v2 protein IDs 2257338, 2257340 and 2257352, respectively) revealed the existence of several CCTX homologues (Tayyrov et al, 2018; Kombrink et al, 2019) over the full length of the protein, specific to the fungal kingdom (Fig. 1B, Fig. S1D; Table S1). Most homologues were detected in Agaricomycetes, with a remarkable expansion in the genus *Pisolithus*. Additionally, a few homologues were encoded by Ascomycetes.

**Figure 1.**
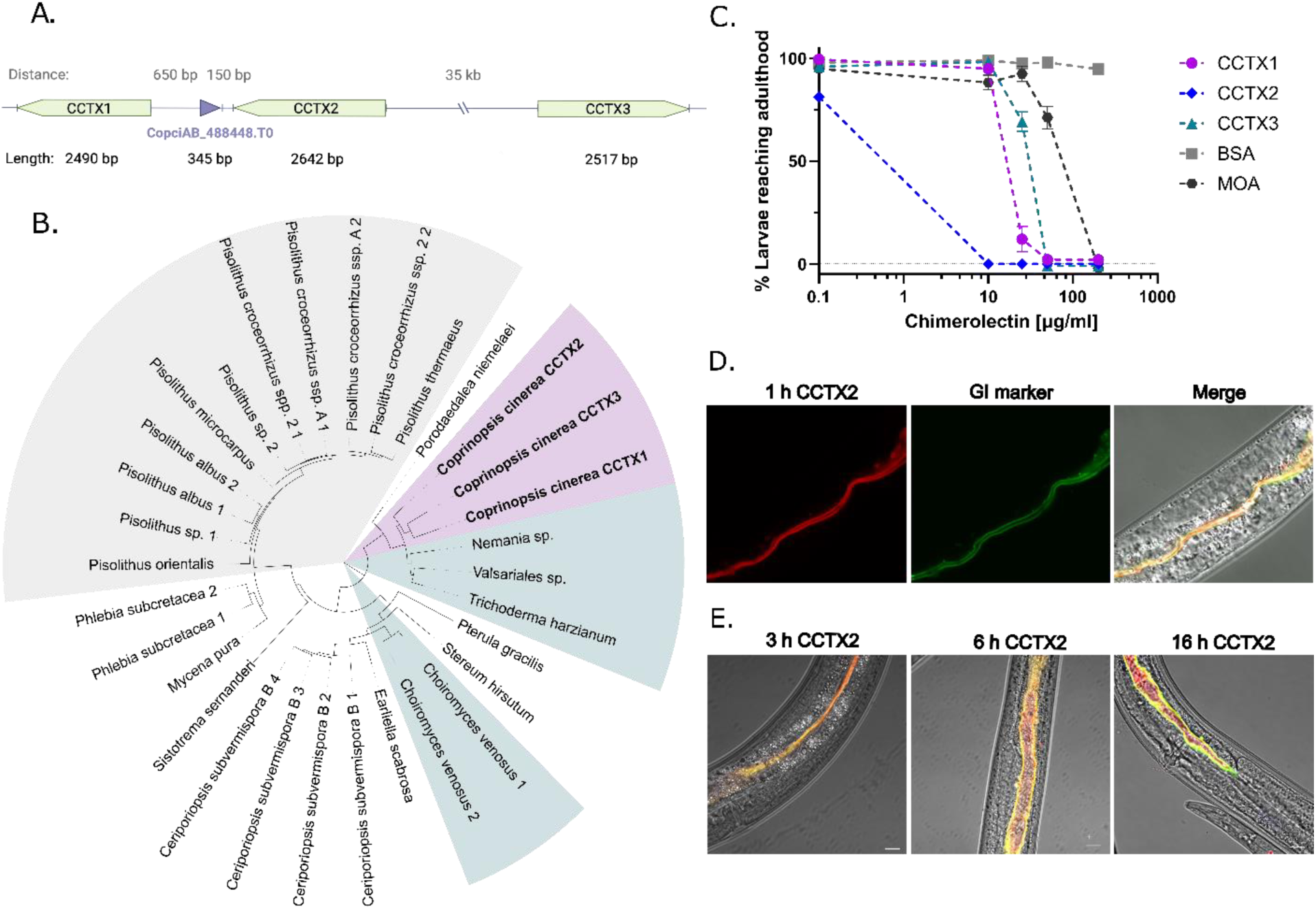
The nematotoxin CCTX2 is a member of a large family of fungal chimerolectins. **A.** Location of CCTX paralogs in the *C. cinerea* genome (*Coprinopsis cinerea* AmutBmut v.2 annotation from JGI). **B.** Phylogenetic analysis of full-length homologues of CCTX2. Paralogues in *C. cinerea* are highlighted in pink, homologues in *Pisolithus* spp. in gray and homologues in Ascomycetes in turquoise. See Table S1 for the JGI protein IDs of all displayed homologues and Fig. S1A for a sequence alignment of homologues from the monophyletic clade including CCTX1-3. **C.** Nematotoxicity assays of the paralogous CCTX1, CCTX2 and CCTX3. Development of *C. elegans* N2 larvae was arrested in L1 larvae in the presence of either of the three proteins at concentrations >10 μg/ml. The MOA chimerolectin was included as positive control, BSA served as negative control. Data points with error bars indicate means of N = 4 biological replicates with standard error of the mean (SEM). Overlapping data points were nudged by +/− 2 data units in Y for better readability. **D** and **E.** Localization of CCTX2-TAMRA upon feeding to L4 larvae of *C. elegans* strain GK70. This strain expresses PGP-1-GFP as intestinal apical membrane marker. Images in panel D were taken 1 h after feeding with CCTX2-TAMRA. **E.** Time course of intoxification by CCTX2. After 3h, feeding of CCTX2-TAMRA leads to aberrant dilatation of the intestinal lumen. Pictures were taken at the rear end of the animal. Scale bar = 10 µm.

The specific induction of *cctx1* and *cctx2* by *A. avenae* suggests a function of these genes in the defense of *C. cinerea* against fungivorous nematodes. Indeed, we previously showed that CCTX2 is toxic towards the nematode model species, *Caenorhabditis elegans* (Plaza et al, 2016). To test if all three paralogous genes encode defense effector proteins, we produced the proteins in *E. coli* and tested their toxicity against *C. elegans.* All three paralogous proteins inhibited the development of *C. elegans* L1 larvae at low concentrations (<50 µg/ml, Fig. 1C). The inhibition of larval development was comparable to the effects of the *Marasmius oreades* chimerolectin MOA and much stronger than for the *C. cinerea* hololectin CCL2 (Fig. S2) (Wohlschlager et al, 2011). Feeding of TAMRA-labeled CCTX2 protein to L4 staged *C. elegans* GK70 larvae (harboring GFP-labeled apical plasma membrane marker protein PGP-1) revealed colocalization of the chimerolectin CCTX2 and PGP-1 with the intestinal epithelium (Fig. 1D). Accordingly, exposure to CCTX2 resulted in clear morphological changes of the intestine, such as gross distension of the intestinal lumen and an increasingly undulating phenotype of the apical plasma membrane after more than 2h of feeding (Fig. 1E). These morphological changes increased in severity over time, leading to death of the worms around 16 to 24 h after toxin administration (data not shown). Similar morphological changes to the intestinal phenotypes were previously described for other nematotoxic fungal lectins, such as the chimerolectin MOA and the two *C. cinerea* hololectins CGL2 and CCL2 (Butschi et al, 2010; Wohlschlager et al, 2011; Schubert et al, 2012; Stutz et al, 2015). Besides *C. elegans*, we found that CCTX2 was also toxic to the larvae of two other *Caenorhabditis* species (Fig. S1C). A similarly broad specificity as CCTX2 was also observed for the chimerolectin MOA but not for the hololectin CGL2x.

### CCTX2 fold comprises four β-trefoil domains and a C-terminal domain of unknown function (DUF)

Sequence analysis revealed that CCTX1, CCTX2 and CCTX3 exhibit four ricin B chain-like domains at the N-terminus, followed by a 220-230 amino acid domain with no homologue in the known proteome (Fig. S1D). The first two domains of each orthologue contain the canonical (QxW)_3_ motif, characteristic of R-type lectins (Fig. S1D) (Yao et al, 2011), implying a possible lectin function (*i.e.*, binding to glycans or glycoconjugates). An *in silico* structural model, generated in pre-AlphaFold era (Jumper et al, 2021) using the Robetta server (Kim et al, 2004; Yang et al, 2020), showed that the ricin B chain-like domains adopt the stereotypical β-trefoil fold (BTF); no fully reliable model could be generated for the C-terminal domain (Fig. S3). To confirm these predictions and gain a better understanding of the molecular function of the individual domains and their interplay, we determined the CCTX2 structure by single-particle cryo-EM to a maximum resolution of 3.2 Å (Fig. 2, Fig. S4; Table S2). Residues 1 to 787 (full sequence) were visible in the reconstructed volume; regions with poorly resolved density are reported in Fig. S3D and Table S3. The structure shows a single polypeptide chain with five identifiable domains connected by short linkers, matching extant predictions (Fig. 2A); domain boundaries are reported in Table S4. Overall, the protein adopts a compact, globular fold (Fig. 2A and 2B). *B*-factor analysis shows the presence of two distinct structure-functional units: BTF domains 1-3, characterized by low flexibility (low *B*-factors), and domain 4-5, showing much higher flexibility/disorder (Fig. 2C). The four ricin-B-chain-like domains adopt the expected β-trefoil fold (BTF) and are arranged in a rhomboid shape, forming a cradle that supports the fifth domain (Fig. 2B). Each BTF domain displays the characteristic arrangement of the α, β and γ subdomains around a pseudo-threefold axis (Fig. S5A); α, β and γ subdomain boundaries are reported in Table S4. A per-domain search for structural homologues using the DALI server (Holm, 2022) returned matches among other fungal lectins and bacterial toxins (Table S6). Structural alignment shows strong similarity among the four BTF domains, despite low sequence conservation (Table S5); however, a visual inspection revealed that the fourth BTF domain contains an incomplete β subdomain (Fig. S5A). The putative sugar-binding sites in all the BTF domains of CCTX2 are located on the outer side of the ‘cradle’ and are thus fully exposed to the solvent (Fig. S5B and S5C). The reconstructed Coulomb potential map showed additional density (Fig. S5D) at the sugar binding pocket of subdomains β and γ of the first BTF domain and subdomain α of the second BTF domain (sites marked in magenta in Fig. 2A and S5). The detected density can probably be traced to residual D-galactose, used throughout the purification process. The canonical (QxW)_3_ motifs of R-type lectins are usually involved in the binding of glycans (Yao et al, 2011), which is in good agreement with the presence of an intact (QxW)_3_ motif only in the first two BTF domains. Finally, three Cys residues (514, 515, 550) form a putative Zn^2+^ binding site, well nestled into BTF domain 4 (Fig. 2B, inset).

**Figure 2.**
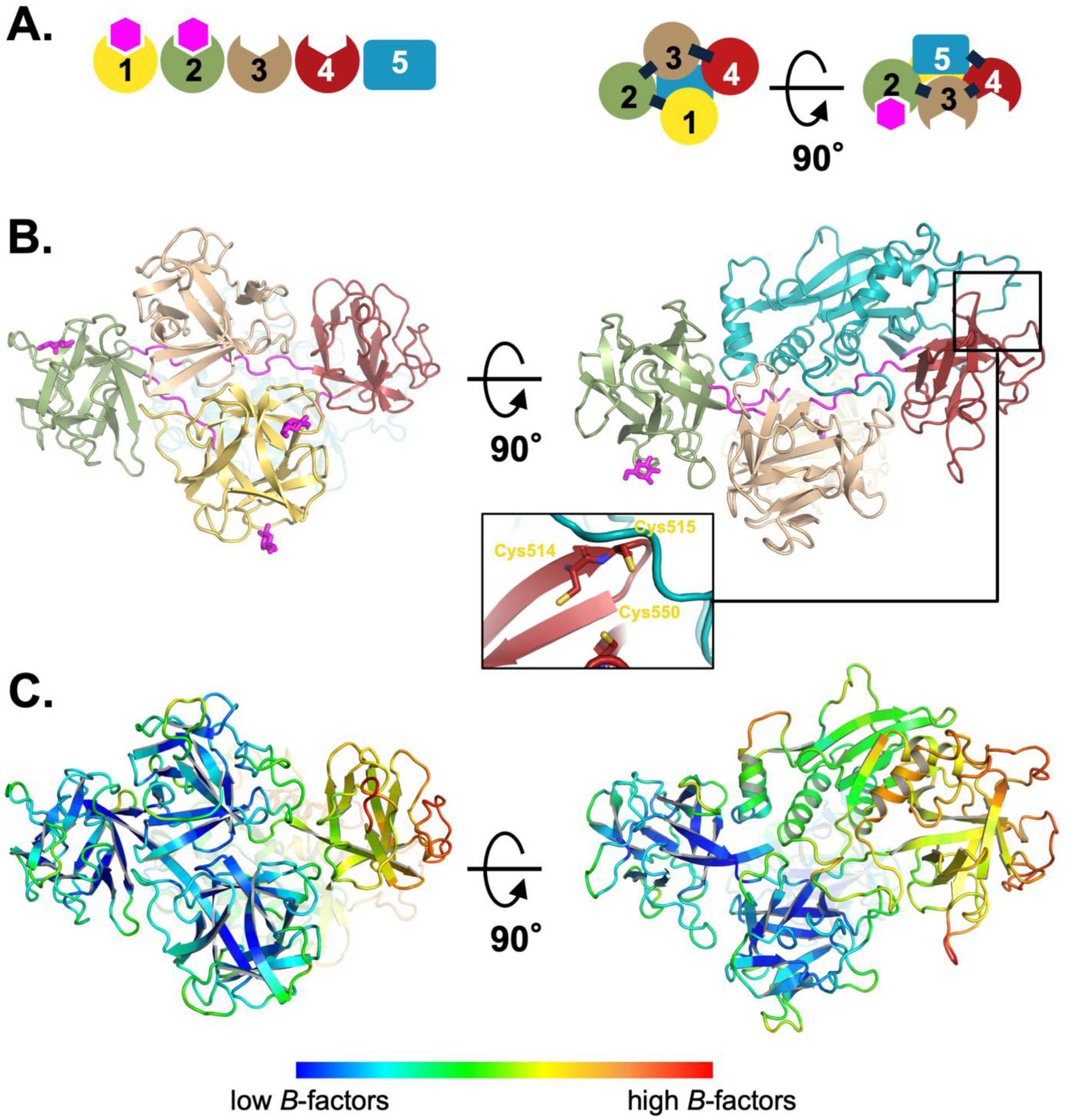
Cryo-EM structure of CCTX2. **A.** Schematic diagram of the domains of CCTX2 and their topology in the protein tertiary structure. Each domain is represented by a different color to be easily located in the folded protein: yellow (β-trefoil domain/BTF domain 1), green (BTF domain 2), wheat (BTF domain 3), red (BTF domain 4) and teal (C-terminal domain of unknown function; DUF). The hexagons in magenta represent sugars observed bound to the BTF domains with proposed lectin activity (see also panel B and Fig. S5). **B**. Cartoon representation of the CCTX2 structure. The domains follow the color code assigned in A, with linkers between domains colored in magenta. D-galactose molecules (see Fig. S5)—likely remnants from the purification procedure—are depicted as magenta-colored sticks. They represent the Coulomb potential density at the β and γ sugar binding sites of domain 1 and the α sugar-binding site of domain 2. The inset shows the position of a putative zinc-binding site **C.** Schematic diagram of CCTX2, colored by *B*-factors of Cα atoms, according to the color chart on the bottom.

The fifth domain (amino acids 557 to 787) adopts a unique α+β fold (Fig. 3), bearing no similarity to any known protein structure. A search for structural homologues was performed using the DALI server (Holm, 2022). Either the full domain or its core portion (residues 557-570 + 620-655 + 684-745) did not return any significant hit, indicating the discovery of a new α+β-type protein fold. The core of the C-terminal domain is formed by a seven-stranded antiparallel β-sheet, supported by two α-helices; the β-sheet changes orientation between strands 6 and 7 (Fig. 3A and 3B). A large portion of the β-sheet is solvent-exposed and forms a potential catalytic cleft. The core of the domain is decorated with two large insertions. The first insert (residues 571-619) contains a helix and a long stretch of random coil that tightly interact with the third BTF domain; the second insert (residues 656-683) forms a two-helix subdomain, packed at the sides of the central core (Fig. 3B and 3C). The C-terminal residues (746-787) form a long random coil segment, circling the fourth BTF domain and ending in a helix and another random coil, sandwiched between the first insert and the central core (Fig. 3B and 3C). Interestingly, the C-terminal loop is in direct contact with the putative Zn^2+^ binding site of CCTX2 (Fig. 2B, inset).

**Figure 3.**
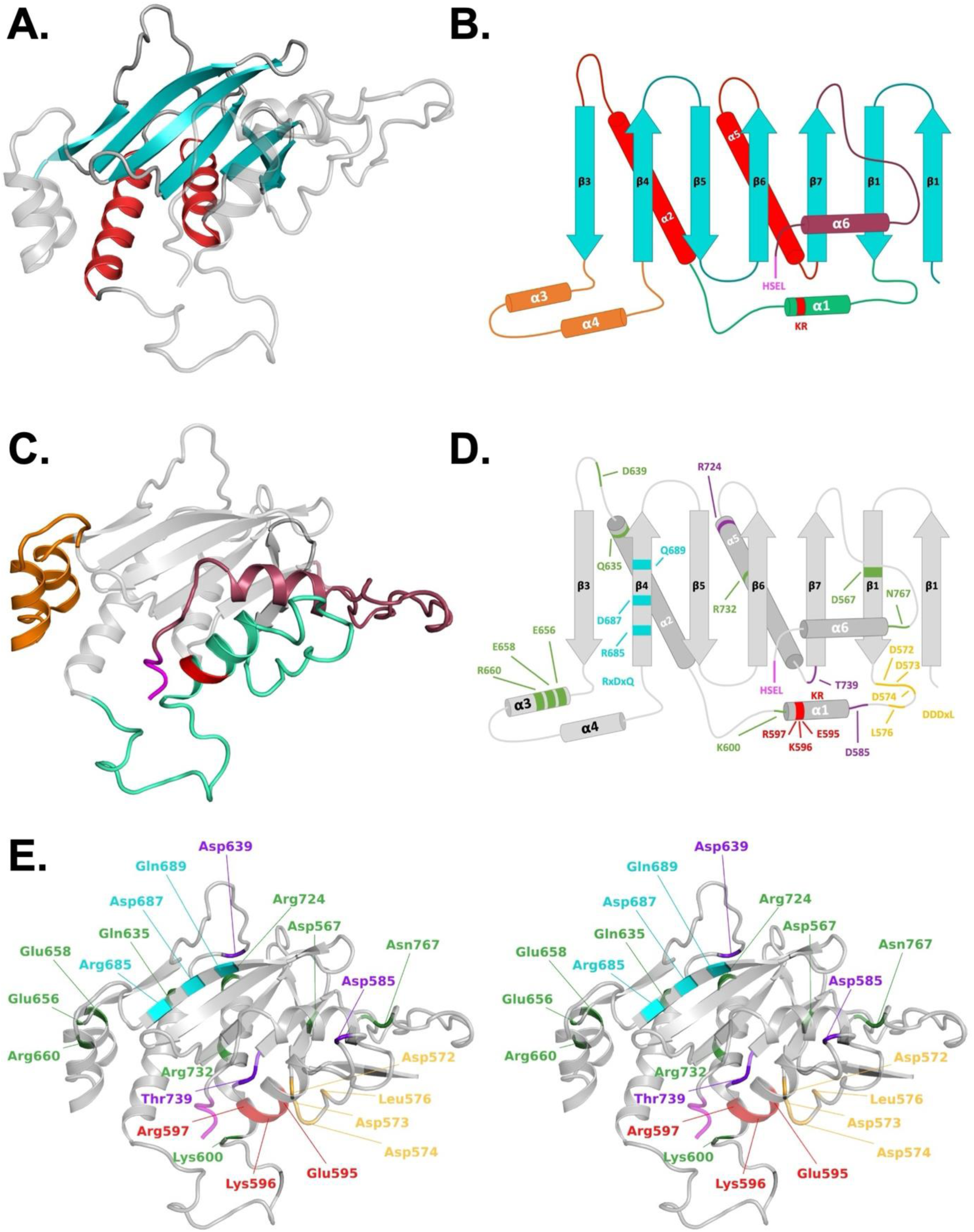
CryoEM structure of CCTX2 C-terminal Domain of Unknown Function (DUF). **A.** Cartoon representation of the C-cryo-EM structure of the C-terminal DUF. The core elements of the fold, the central seven-stranded β-sheet and the two supporting α-helices are indicated in teal and red, respectively. **B.** Topology diagram of the C-terminal DUF indicating insertions and conserved sequons. The KR sequon for kexin cleavage (residues 596-597) in the first insertion (residues 571-619, light green) is highlighted in red. The two-helix insertion (residues 656-683) is shown in orange. The C-terminus (residues 746-783) is colored dark red, with the HSEL sequon (residues 784-787) highlighted in magenta. **C**. Cartoon representation of the DUF fold indicating the same insertions and conserved sequons as in panel B. **D.** Topology diagram of the C-terminal DUF reporting the position of amino acids targeted by site mutagenesis. **E.** Stereo cartoon representation of the C-terminal DUF, with residues targeted by Ala-scanning substitutions (green) mapped onto the cryo-EM structure. Positions corresponding to the key functional motifs KR (Glu595, Lys596, Arg597; red), RxDxQ (Arg685, Asp687, Gln689; cyan), DDDxL (Asp572, Asp573, Asp574, Leu576; yellow), HSEL (C-terminal residues 784-787; magenta) and three variants significantly affecting toxicity (Asp585, Arg724, Thr739; purple) have been marked in different colors.

### Nematotoxicity of CCTX2 is dependent on the two N-terminal lectin domains and the C-terminal DUF domain

To characterize the molecular functions of the different CCTX2 domains, we generated both N- and C-terminally truncated variants of the protein (Fig. 4A). Since both sequence and structural analysis predicted a lectin function for the first two BTF domains, we produced a variant lacking these two domains (residues 304 to 787), termed CCTX2ΔN. To assess carbohydrate-binding, we probed AlexaFluor488-labeled versions of both full-length CCTX2 and CCTX2ΔN for binding to the CFG mammalian glycan array v5.1, using the service provided by the Consortium of Functional Glycomics (CFG, Emory University, USA). The data show that CCTX2 is indeed a lectin with specificity towards LacNAc(Galβ1-4GlcNAc)- and LacdiNAc(GalNAcβ1-4GlcNAc)-containing glycans (Fig. S6 and supplementary data files). No binding was detected for CCTX2ΔN (Supplementary data files), but the protein was soluble suggesting structural and functional integrity. Overall, results indicate that the first two BTF domains of CCTX2 are required for its carbohydrate-binding activity, in agreement with the sequence predictions and the structural results. To test whether the carbohydrate-binding activity of CCTX2 is necessary for its nematotoxicity, we analyzed CCTX2ΔN with our nematotoxicity assay (Fig. 4B). We found that this protein variant was not nematotoxic, strongly suggesting that the lectin function is essential for the nematotoxicity of CCTX2.

**Figure 4.**
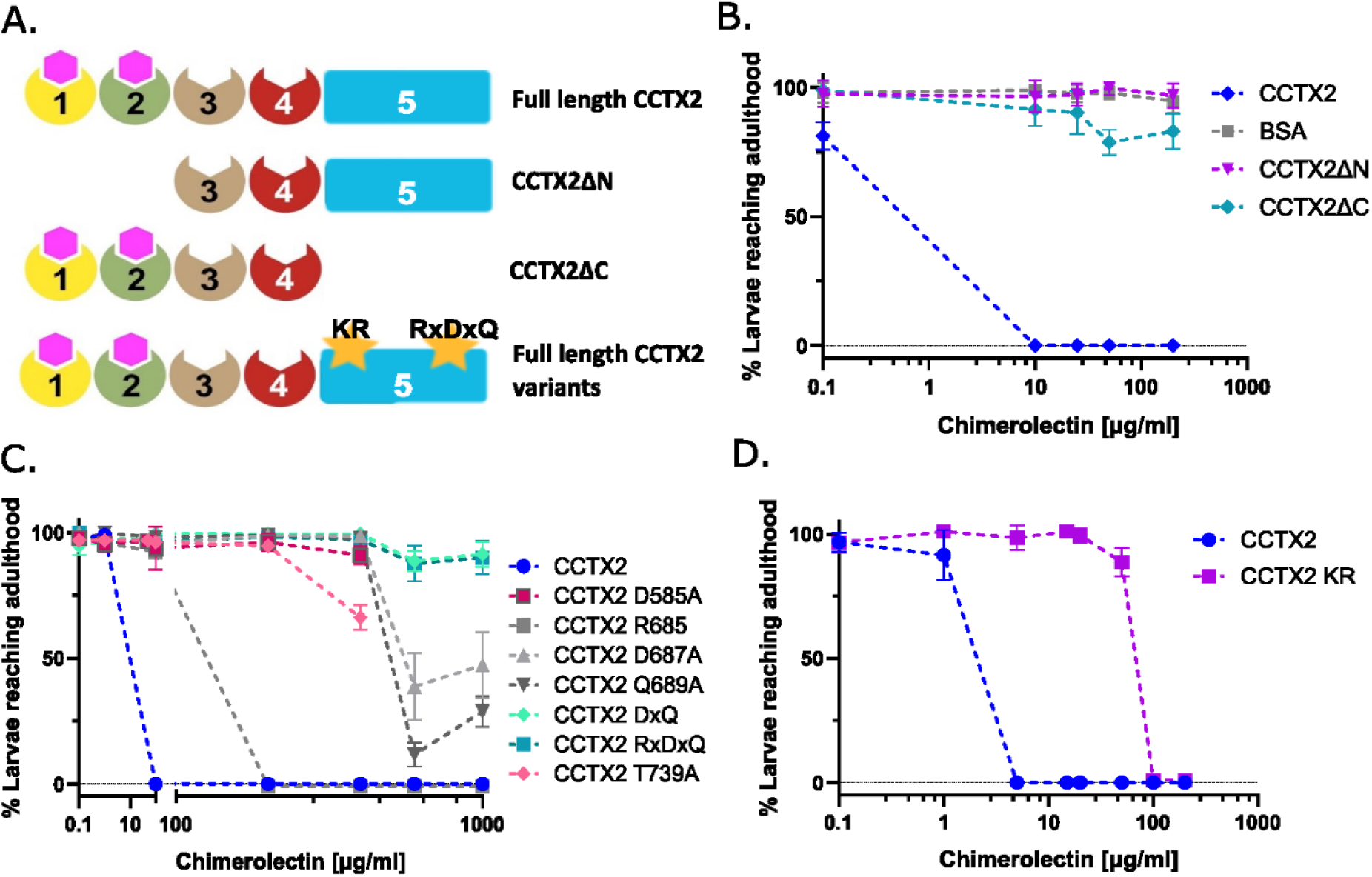
The two N-terminal BTF lectin domains and the C-terminal DUF domain are required for CCTX2 nematotoxicity. **A.** Schematic representation of the CCTX2 architecture with KR and RxDxQ motifs indicated by yellow stars (see main text for details). **B.** *C. elegans* nematotoxicity assay with full-length CCTX2 (blue), a CCTX2 variant lacking the two N-terminal BTF domains (CCTX2ΔN, magenta) and a variant lacking the C-terminal domain of unknown function (CCTX2ΔC, teal), with BSA used as negative control. **C**. *C. elegans* nematotoxicity assay with CCTX2 variants in the RxDxQ motif either as single substitutions (grey, R685A, D687A and Q689A) or double (DxQ, light teal) or triple alanine replacements (RxDxQ, teal). Point mutations D585A and T739A also conferred resistance to the toxin. **D.** Resistance of *C. elegans* to CCTX2-KR (E595A, K596A, R597A) in nematotoxicity assay. Data points with error bars indicate means of N = 4 biological replicates with standard error of the mean (SEM). Overlapping data points were nudged by +/− 2 data units in Y for better readability.

Although the C-terminal domain does not show any structural similarity to known proteins, it is well conserved among the three CCTX paralogues, with a sequence identity of approximately 40% (residues 555 to 787). To assess its role for nematotoxicity, we created a variant of CCTX2, termed CCTX2ΔC, which lacks this domain (residues 570 to 787) (Fig. 4A). This variant exhibited severely reduced nematotoxicity, indicating that this domain is also essential to mediate fungal defense (Fig. 4B). Currently, we can only speculate about the molecular function of this domain. An amino acid sequence alignment between the members of the monophyletic clade of CCTX homologues revealed several conserved amino acid residues within the domain (Fig. 1B and S1D). We individually replaced sixteen of these residues with alanine and evaluated the toxicity of the resulting CCTX2 variant proteins in our nematotoxicity assay against *C. elegans* (Fig. S1D and Table S7). We also mapped the residues onto the structure (Fig. 3D). Nine of these CCTX2 variants showed a reduction in toxicity of CCTX2 towards *C. elegans* (Table S7, Fig. 4C and S7A and S7B). Of these, substitutions D687A and Q689A had the strongest effects with a reduction of the toxicity of CCTX2 towards *C. elegans* by two orders of magnitude (Fig. 4C). A double replacement of both residues (‘DxQ’ variant), as well as a triple variant (‘RxDxQ’ variant) that includes neighboring residue Arg685, whose individual substitution caused only a weak reduction in toxicity (Fig. 4C), rendered the protein completely non-toxic. We renamed these three amino acids the ‘RxDxQ motif’ (referring to Arg685, Asp687 and Gln689 in CCTX2) (Fig. 4C). In the CCTX2 structure, this motif is exposed on the surface of the central β-sheet (Fig. 3D). The single substitutions D585A and T739A in the C-terminal domain also resulted in a significant reduction in toxicity (non-toxic up to a concentration of 500 mg/l), while the variants R724A as well as Q635A and E656A were less strongly reduced, being non-toxic at a concentration below 200 mg/l and 100 mg/l, respectively (Fig. S7A, Fig. S8 and Table S7).

Apart from the ‘RxDxQ motif’, we found that the C-terminal domain contains a conserved putative cleavage site (Lys596 and Arg597 in CCTX2) for the Golgi-localized endoproteinase Kexin (Fig. 3B-E). Substitution of these two residues (and the preceding Glu595) to alanine in CCTX2 resulted in a reduction of toxicity of this protein variant, CCTX2-KR, against *C. elegans* (Fig. 4D). This site is embedded in the second structural insert, on a solvent-exposed α-helix (Fig. 3C). To check whether this motif was functional, purified CCTX2 protein was incubated with Kex2p protease from *Saccharomyces cerevisiae* and the reaction was subjected to a subsequent pulldown experiment using Sepharose beads. Analysis of the various fractions by SDS-PAGE and immunoblotting, using a custom-made anti-CCTX2 antiserum, showed that CCTX2 was cleaved into two fragments of ≈23 and ≈67 kDa, corresponding to the molecular masses expected from a proteolytic processing at the presumptive cleavage site (Fig. S7C). Intriguingly, however, Kex2p cleavage alone was insufficient to separate the two fragments, as we could not detect the C-terminal fragment, neither in the pulldown flow-through nor in the wash. In conclusion, the C-terminal fragment of CCTX2 is tightly associated with the N-terminal fragment and can only be separated under denaturing conditions, such as SDS-PAGE (Fig. S7C).

Finally, we noticed a conserved motif, Asp572 to Leu 576, which we refer to as DDDxL, located in between BTF4 and the ‘KR motif’ (Fig. 3D-E). Substitution of all three Asp residues and the terminal Leu residue of this motif to alanine resulted in a protein (CCTX2-DDDxL) that was reduced in toxicity (non-toxic at a concentration below 100 mg/l) (Fig. S7A, Fig. S8 and Table S7) to similar extent as the single substitutions of residues Gln635 and Glu656 (see above).

Taken together, this mutational analysis confirmed that the C-terminus of CCTX2 is essential for its nematotoxicity but did not pinpoint the biochemical function of this domain.

### CCTX2 binds to C. elegans intestinal epithelial glycosphingolipids in vivo and in vitro

The dependence of the nematotoxic activity of CCTX2 on two lectin domains (Fig. 4B) and the localization of TAMRA-labeled CCTX2 to the intestinal epithelial membrane in *C. elegans* (Fig. 1C and 1D) suggest that this chimerolectin targets a glycoconjugate present in the apical intestinal plasma membrane of nematode enterocytes. The specificity of CCTX2 towards LacNAc, and LacdiNAc containing motifs (Fig. S6), the binding of nematotoxic chimerolectin MOA to *C. elegans* glycosphingolipids (GSLs) (Wohlschlager et al, 2011) and the presence of LacdiNAc in the core of the glycan part of *C. elegans* GSLs (Barrows et al, 2006) point to these glycoconjugates as putative target molecules. The biosynthesis of the *C. elegans* GSL glycans occurs by the sequential action of the enzymes BRE-3 (β1,4-mannosyltransferase), BRE-5 (β1,3-*N*-acetylglucosaminyl transferase), BRE-4 (β1, 4 *N*-acetylgalactosaminyl transferase) and BRE-2 (β1,3-galactosyltransferase) (Fig. 5A) (Barrows et al, 2006). To test the role of GSLs in toxin binding, loss-of-function *C. elegans* mutants of the four GSL glycosyltransferase-encoding genes were tested for resistance towards CCTX2. All the tested *C. elegans* GSL mutant strains displayed increased resistance to CCTX2, except for the *bre-2(ye31)*-mutant strain (Fig. 5B). This result strongly suggests that the LacdiNAc motif in the core of the *C. elegans* GSLs is the glycoepitope targeted by CCTX2. To demonstrate direct binding of CCTX2 to *C. elegans* GSLs, we performed CCTX2 protein overlays for GSLs isolated from wildtype and mutant *C. elegans* strains. *C. elegans* GSLs were separated by TLC and overlayed with biotinylated CCTX2, using biotinylated MOA as positive control. The results showed binding of CCTX2 to GSLs from *C. elegans* N2 wild-type and the *bre-2(ye31)* mutant strain, similar to the MOA positive control (Fig. 5C). No binding of CCTX2 or the MOA positive control to the GSLs isolated from *bre-3, -4 or -5* strains was observed, consistent with the resistance of those strains to both proteins (Wohlschlager et al, 2011). In addition, we could not observe any binding to GSLs for the CCTX2ΔN variant, which lacks the carbohydrate-binding domains (Fig. 5C).

**Figure 5.**
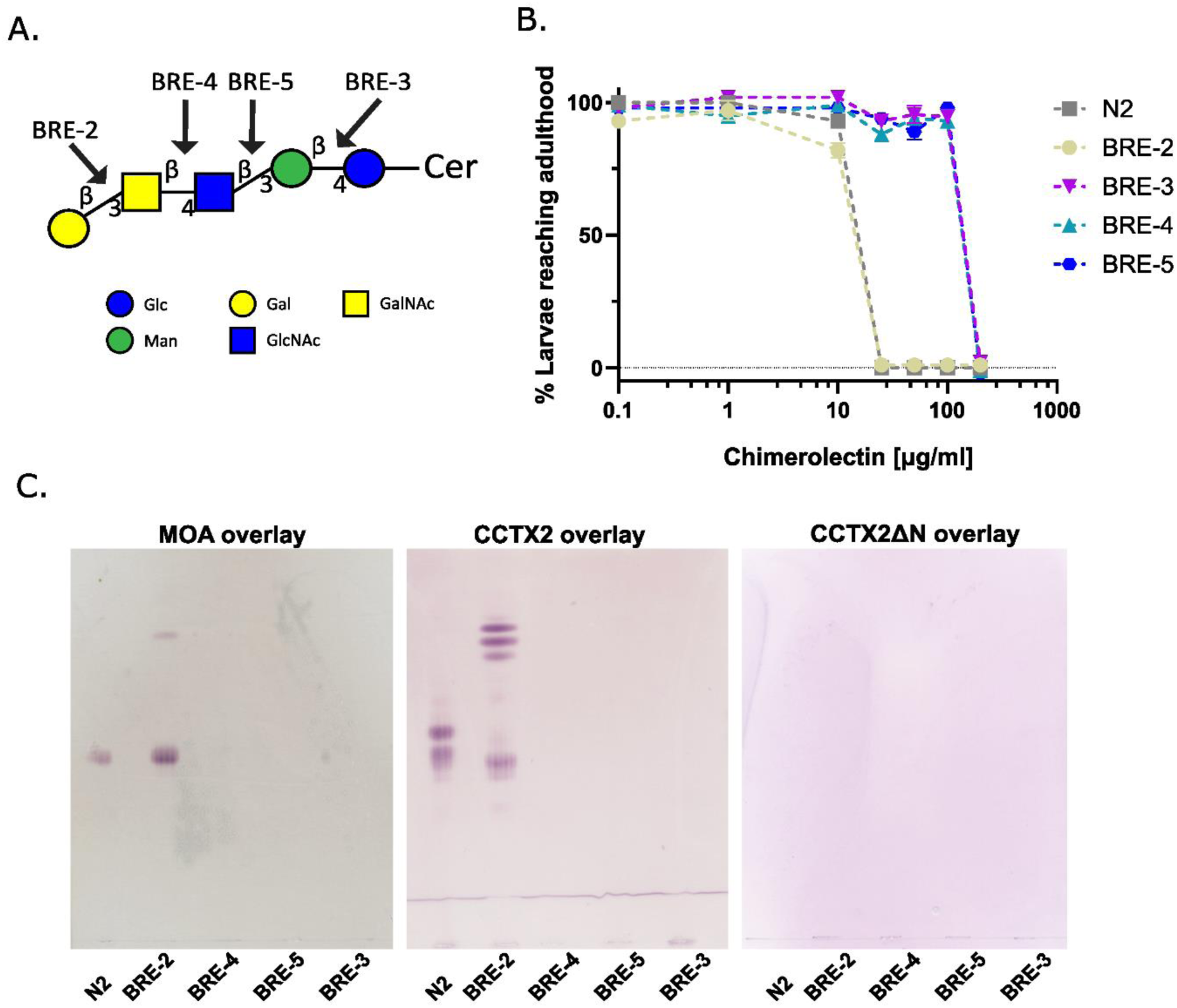
CCTX2 binds to *C. elegans* intestinal epithelial glycosphingolipids *in vivo* and *in vitro*. **A.** Schematic representation of the arthroseries GSL core including the *C. elegans* enzymes responsible for the addition of individual monosaccharides. The ceramide-linked core is synthesized by sequential action of the enzymes BRE-3, BRE-5 and BRE-4; GalNAc is transferred by BRE-2. Monosaccharides are represented as introduced by the Consortium for Functional Glycomics (CFG) (Varki et al, 2015). **B.** Toxicity assay on *C. elegans* GSL biosynthesis mutants *bre-3(ye26), bre-5(ye-17), bre-4(ye27)* and *bre-2(ye31).* L1 larvae of mutant and wild-type worms were fed with different concentrations of CCTX2 (up to 200 µg/ml) and scored for the percentage of larvae reaching the L4 stage/adulthood. *C. elegans* N2 and the mutant for the GalNAc transferase *bre-2(ye31)* were sensitive to CCTX2 at low concentrations. Data points with error bars indicate means of N = 4 biological replicates with standard error of the mean (SEM). Overlapping data points were nudged by +/− 2 data units in Y for better readability. **C.** TLC overlays of upper phase GSLs isolated from the respective *C. elegans* GSL biosynthesis mutants and N2 wild-type strains, blotted with 150 nM of the indicated biotinylated protein.

### CCTX2 undergoes endocytosis and retrograde trafficking in C. elegans enterocytes

Many bacterial GSL-binding AB toxins (*e.g.,* Shiga or cholera toxins (Sandvig et al, 2014)), are endocytosed upon binding to a glycolipid receptor (Fig. 6A). We hypothesize therefore that the fungal chimerolectins CCTX2 and MOA follow a similar pathway. In contrast, toxic hololectins, which bind to and cluster *N*-glycoprotein receptors, *i.e.* protein-bound glycans (*e.g.,* CGL2 (Butschi et al, 2010) and CCL2 (Schubert et al, 2012)), might not require internalization and exert toxicity by a different mechanism (Stutz et al, 2015; Bleuler-Martinez et al, 2017). To assess if endocytosis of CCTX2 was required for its nematotoxicity, we fed TAMRA-labeled CCTX2 to L4 larvae of *C. elegans* GK70, a strain carrying the apical plasma membrane marker PGP-1 fused to GFP and monitored the localization of CCTX2 using confocal fluorescence microscopy. The fungal chimerolectin MOA and the hololectin CCL2 were included for comparison. After 3h of feeding, intracellular TAMRA-positive vesicles were observed in *C. elegans* enterocytes (Fig. 6B). Several endocytic vesicles co-stained with PGP-1::GFP were detected, indicating active endocytosis of the chimerolectin CCTX2. Indeed, we also observed such endocytic vesicles for TAMRA-labeled and GSL-binding chimerolectin MOA. In contrast, we did not detect TAMRA-CCL2 in endocytic vesicles, which is in agreement with previous studies (Stutz et al, 2015). Additionally, we could not detect endocytic vesicles with the CCTX2ΔN variant (Fig. 6B), suggesting again that GSL receptor binding is necessary for endocytosis of the protein toxin.

**Figure 6.**
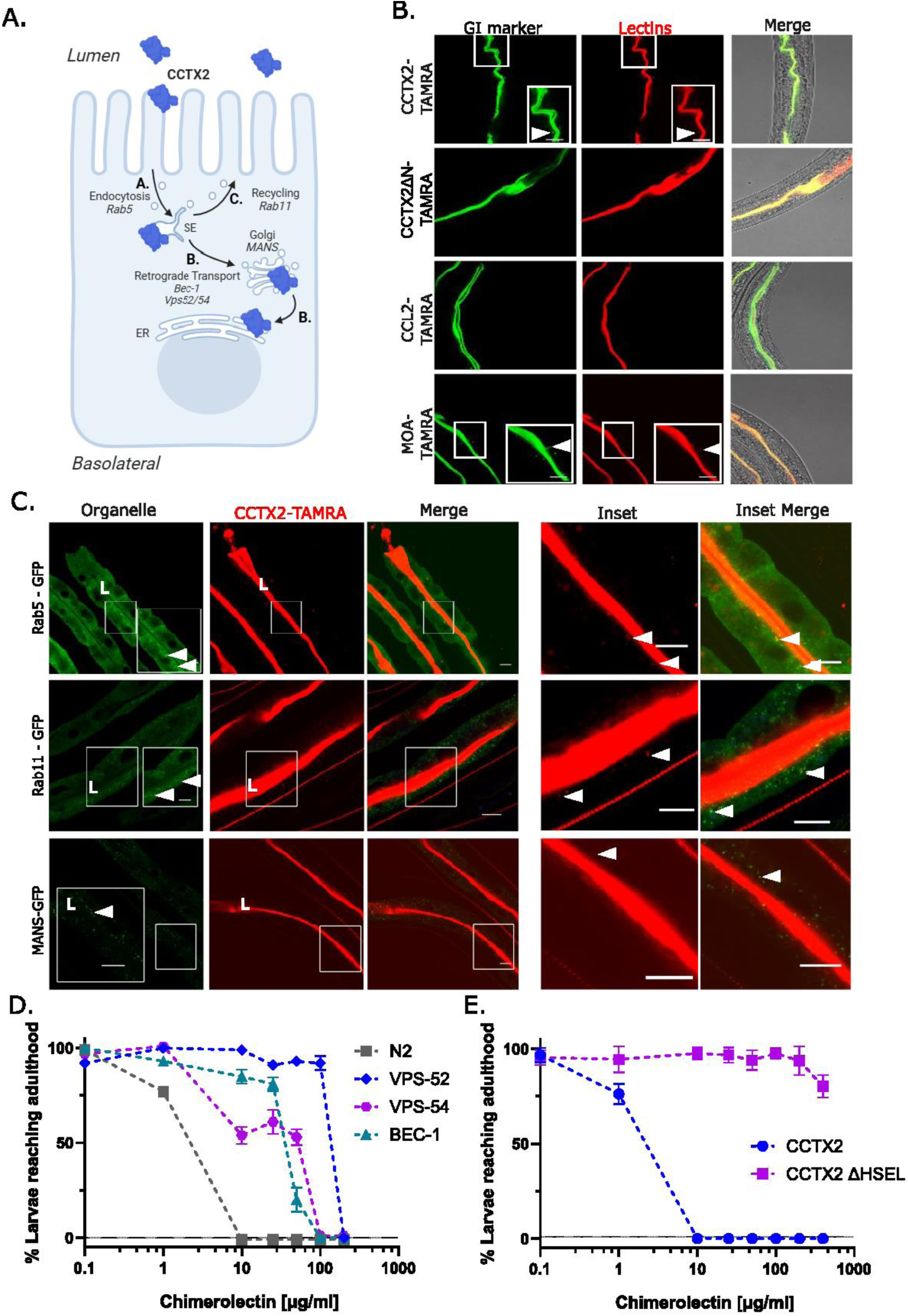
Endocytosis and retrograde trafficking of CCTX2 in *C. elegans* enterocytes. **A.** Schematic representation of the endocytic network in *C. elegans* enterocytes. After endocytosis and fusion of the endocytic vesicles with the sorting endosome (SE) (A.), the cargo is sorted either for recycling back to the plasma membrane (C.) or for retrograde trafficking to the trans Golgi network (TGN), Golgi and endoplasmic reticulum (ER) (B.). Cargo destined for degradation is sorted into the late endosome (LE) and the lysosomal pathway. **B.** Internalization of CCTX2 into *C. elegans* enterocytes. Fluorescently-labeled CCTX2 (red), truncated CCTX2ΔN (red), the GSL-binding chimerolectin MOA (red) or the N-glycan binding lectin CCL2 (red) were fed to *C. elegans* GK70 (green) for 3 h at 20°C. Arrowheads indicate fluorescent puncta (vesicles) as a sign of endocytosis. These vesicles appear red (due to the TAMRA-labelled cargo), green (due to the intestinal apical membrane marker PGP-1::GFP) or yellow (merge). **C.** Co-localization of CCTX2 with endocytic markers. *C. elegans* RAB-5::GFP, RAB-11::GFP as well as the Golgi marker α-Mannosidase II (MANS::GFP) were used as endocytic and retrograde trafficking markers to test for co-localization with CCTX2. TAMRA-labeled CCTX2 was fed to the respective *C. elegans* L4 larvae for 6 h at 20°C and monitored for co-localization. Images show the intestinal cross-sections. Arrowheads indicate overlapping puncta (vesicles). ‘L’ indicates the position of the intestinal lumen. Scale bars = 10 µm. **D.** Nematotoxicity assay with CCTX2 with different *C. elegans* retrograde trafficking mutants: VPS-52 (VC625) and VPS-54 (VC985) bear mutations in the GARP complex subunits VPS-52 and VPS-54, respectively. BEC-1 is required for retromer localization to endosomes. VPS-36 (VC947) carries mutation in the ESCRT II subunit. **E.** Nematotoxicity assay of CCTX2 variant lacking the terminal HSEL sequon. Data points with error bars indicate means of N = 4 biological replicates with standard error of the mean (SEM). Overlapping data points were nudged by +/− 2 data units in Y for better readability.

To confirm the endocytosis of CCTX2 in enterocytes, we used *C. elegans* strains with fluorescently marked endocytic compartments (Fig. 6A and 6C). For this purpose, we fed TAMRA-labeled CCTX2 to *C. elegans* RAB-5::GFP (Rab5 is a small GTPase present on early endosomes (Bucci et al, 1992)), RAB-11::GFP (Rab11 is a small GTPase on recycling endosomes (Hoekstra et al, 2004; Jing & Prekeris, 2009; Sato et al, 2014)); and MANS::GFP (α-Mannosidase II, an enzyme localized in the Golgi (Shah et al, 2008)) strains. After 6h of feeding double positive intracellular vesicles containing both, CCTX2 and the endocytic marker protein were observed in enterocytes of the RAB-5::GFP strain (Fig. 6C), confirming that fluorescently labeled CCTX2 was endocytosed upon receptor binding. Colocalization of CCTX2 and RAB-11::GFP was also observed (Fig. 6B). Localization of CCTX2 to downstream organelles such as the Golgi apparatus (*C. elegans* MANS::GFP) was rarely detected.

After being endocytosed, many GSL-binding AB toxins hijack retrograde vesicle trafficking to reach the endoplasmic reticulum (ER) (Teter, 2013). From this compartment, many AB toxins are released into the cytoplasm by retro-translocation, where they exert their toxic effect (Sandvig & van Deurs, 2005; Sandvig et al, 2010; Sandvig et al, 2013). The retrograde trafficking of CCTX2-containing vesicles and its requirement for nematotoxicity was evaluated using *C. elegans* strains deficient in different components of the retrograde trafficking machinery. Deletions in the GARP (Golgi-associated retrograde protein) complex subunits VPS-52 and VPS-54 rendered the *C. elegans* partially resistant to intoxication with CCTX2 (Fig. 6D). In addition, a heterozygous knock-out mutant for an ESCRT-II subunit, and the VPS-34-recruiting protein BEC-1 mutant led to increased resistance to CCTX2 compared to the *C. elegans* N2 wild-type strain (Fig. 6D). Neither of these strains showed increased resistance towards MOA or CCL2, indicating that their resistance towards CCTX2 was toxin-specific and not a pleiotropic effect due to overall changes in membrane or trafficking homeostasis (Fig. S2). Given that the intracellular target for the chimerolectin MOA is unknown, we speculate that this chimerolectin might not require retrograde trafficking to reach its target or uses a different trafficking route.

The three *C. cinerea* paralogues contain a conserved C-terminal four-residue sequence (HSEL) at their C-termini, resembling the canonical ER-retention signal (KDEL) present e.g. in cholera toxin for retention in the ER (Fig. S1D, Fig. 3B and 3C). Removal of the sequon from CCTX2 (CCTX2ΔHSEL) abolished toxicity of the protein towards *C. elegans* N2 (Fig. 6E). This indicates that the presence of an intact HSEL sequon is a prerequisite for toxicity, and its removal likely affects either translocation into the ER or retention of the protein in this compartment.

Taken together, the results suggest that CCTX2 is internalized into early endosomes (Rab5-dependent). From there, the toxin can be recycled back to the plasma membrane via Rab11-positive vesicles or undergoes retrograde trafficking to the TGN, Golgi and the ER facilitated by the GARP complex (VPS-52 and VPS-54) and the ESCRT machinery (BEC-1).

## DISCUSSION

Fungal lectins have been implicated in defense against foraging nematodes or insects (Bleuler-Martínez et al, 2011; Wohlschlager et al, 2011; Tayyrov et al, 2018). In the present work, we describe CCTX1, CCTX2 and CCTX3, three nematotoxic chimerolectins from the basidiomycete *C. cinerea*. We performed an in-depth investigation of CCTX2, the founding member of this new family of fungal chimerolectins, and provide a structure-function analysis by characterizing its cryo-EM structure, its target glycoconjugate and its internalization route in the model nematode *C. elegans*.

### CCTX family of chimerolectins and CCTX2—its holotype

CCTX1, CCTX2 and CCTX3 are the first known representatives of a new family of fungal defense proteins with a mechanism of action reminiscent of bacterial AB toxins. AB toxins are relatively rare in the kingdom of fungi, with MOA and its homologue PSL1a being the most prominent members described so far (Wohlschlager et al, 2011; Juillot et al, 2016). Orthologues of the CCTX family are restricted to the fungal kingdom (Fig. 1B), predominantly in the phyla Basidiomycota and Ascomycota. Intriguingly, the closest sequence homologue of the CCTX toxins can be found in *Trichoderma harzianum*, a pathogen of plants and other fungi without known sexual stage (Gams & Meyer, 1998) (THTX2, UniProt ID: A0A0F9ZAH1).

The *cctx1*, *cctx2* and *cctx3* genes are located next to each other in the genome of *C. cinerea*, suggesting that they are the result of gene duplication events. Similar tandem arrangements of defense genes can be found in *C. cinerea* for nematotoxic hololectins CGL1-CGL2 and CCL1-CCL2 (Boulianne et al, 2000; Schubert et al, 2012) as well as bacteriocidal cysteine-stabilized β-defensin CPP1 (Copsin)/CPP2 (Kombrink et al, 2019). Gene duplications allow organisms to diversify a gene without losing its original function (Moore & Purugganan, 2003). Similar to the other known examples of duplicated *C. cinerea* defense genes, we observed diversification of the *cctx* genes at the level of transcriptional regulation. Whereas *cctx2* is expressed at low level only in the vegetative mycelium and induced upon challenge with fungivorous nematodes (Plaza et al, 2016), *cctx1* and *cctx3* are constitutively expressed at significant levels in young fruiting bodies and vegetative mycelium, respectively (Fig. S1A and S1B).

*In silico* predictions of CCTX2 (Fig. S3A-B) suggested that CCTX family members are composed of five domains: four N-terminal BTF domains and a C-terminal domain of unknown function. This arrangement resembles that of AB_4_ toxins, represented by MTX from *Bacillus sphaericus* and pierisin-1 from *Pieris rapae*, the cabbage butterfly (Treiber et al, 2008; Oda et al, 2017). However, these toxins display a swapped domain organization, with the four BTF domains placed *C-terminally* and the effector domain at the *N-terminus.* The structure of CCTX2, determined by cryo-EM to 3.2 Å resolution, confirms the β-trefoil fold for the N-terminal domains, and reveals their homology with BTF domains from other fungal lectins and bacterial toxins (Table S6), including MOA (Grahn et al, 2007; Grahn et al, 2009), PSL1a (Kadirvelraj et al, 2011) and the *Clostridium botulinum* hemagglutinin component (HA1, (Inoue et al, 2003)). The BTF domains of CCTX2 cluster in a rhomboid fashion, forming a cradle for the fifth domain (Fig. 2A and 2B). This differs from MTX or pierisin-1, where the four BTF domains are arranged in a linear fashion around one side of the effector domain (Treiber et al, 2008) (Fig. S9). Furthermore, the C-terminal domain of CCTX2 does not show any sequence conservation or structural similarity to the effector domain of MTX and pierisin-1, which in both proteins has ADP-ribosyltransferase activity (Treiber et al, 2008; Oda et al, 2017).

BTF domains often have lectin activity, and, accordingly, CCTX2 was confirmed to be a lectin by glycan array analysis, specifically recognizing the LacNAc and LacdiNAc glycoepitopes (Fig. S6). The first two N-terminal BTF domains appear to be required for sugar binding, as the N-terminal deletion construct CCTXΔN failed to bind to the array (Supplementary data files). Although we cannot exclude that the N-terminal deletion causes structural changes that may compromise glycan binding activity, our conclusion is supported by the fact that the protein behaves well upon production and purification and that the putative sugar-binding sites are clustered on the same, solvent-exposed molecular surface (Fig. S5). Moreover, QxW motifs β and γ on the first domain and site α on the second domain show additional density, which likely corresponds to bound D-galactose from the purification process. This indicates that the two N-terminal BTF domains 1 and 2 are important for glycan binding on GSL receptors. The lack of residual density at the putative sugar-binding sites of the third and fourth BTF domains and the less-conserved β subdomain in the fourth BTF domain suggests that BTF domains 3 and 4 have a function different from receptor recognition—potentially for structural support of the C-terminal domain.

### Endocytosis and intracellular trafficking of CCTX2

CCTX2 enters *C. elegans* enterocytes by binding to arthroseries glycosphingolipids carrying the LacdiNAc moiety (Fig. 5). *C. elegans* GSLs are also targeted by the pore-forming nematicidal toxin Cry5B from *Bacillus thuringensis* as well as by the fungal chimerolectin/AB toxin MOA (Griffitts et al, 2005; Wohlschlager et al, 2011). Glycoproteins might provide a secondary entry route, as indicated by the residual toxicity detectable in GSL-deficient mutants *bre-3*, *bre-5* and *bre-4* (Fig. 5B). Receptor binding is likely mediated by the two N-terminal BTF lectin domains, since no GSL binding was detected for CCTXΔN (Fig. 4C). GSL binding triggers internalization of CCTX2, as the toxin colocalized with the apical plasma membrane marker PGP-1 in vesicles within *C. elegans* enterocytes (Fig. 6B). Colocalization of CCTX2 with fluorescent endocytic markers, such as Rab-5 or Rab-11 (*C. elegans* Rab-5::GFP and Rab-11::GFP mutants), was conclusive evidence that CCTX2 undergoes endocytosis (Fig. 6C).

Several AB toxins (*e.g.*, Shiga toxin, cholera toxin, *etc.*) take the same internalization route, followed by retrograde trafficking to the ER and retro-translocation to the cytoplasm (Sandvig & van Deurs, 2005; Cho et al, 2012; Teter, 2013). CCTX2 retrograde trafficking was demonstrated using several *C. elegans* mutants deficient for intracellular trafficking components (Fig. 6/Table S8). These include a mutant in ESCRT-II, mutants for subunits VPS-52 and VPS-54 of the GARP (Golgi associated retrograde protein) complex and a *bec-1* heterozygous mutant (Fig. 6D). In *C. elegans,* BEC-1 mediates autophagy and is required for retrograde transport from endosomes to the TGN (Takacs-Vellai et al, 2005; Ruck et al, 2011). The toxicity resistance of the *bec-1* mutant links the toxic effect of CCTX2 to its trafficking from endosome to the Trans-Golgi-Network (TGN), a necessary step in retrograde trafficking. Moving backward, the GARP complex is required for tethering endosomal-derived vesicles to the Golgi apparatus and is localized within the TGN (Luo et al, 2011). The toxin resistance of the *vps-52* and *vps-54* mutants indicates a strong requirement for TGN transit in CCTX2 trafficking. None of the retrograde trafficking mutants showed resistance to the chimerolectin/AB toxin MOA. This suggests that the observed effects were specific to sorting and trafficking of CCTX2, and not pleiotropic effects (Fig. 6D). Apparently, the retrograde trafficking pathways of CCTX2 and MOA substantially diverge from the endosome onwards. Retrograde trafficking of bacterial AB toxins is often required for toxicity, with ER processing being a fundamental step in toxin activation (Nowakowska-Gołacka et al, 2019). Several bacterial toxins, such as cholera toxin or *Pseudomonas* exotoxin A rely on a KDEL sequon for efficient ER targeting and accumulation by Erd2, the ER protein retention receptor (Lencer & Saslowsky, 2005). CCTX paralogues carry a conserved HSEL sequon at the C-terminus, similar to the ER-retention signal HDEL/KDEL in yeast/mammalian cells. Removal of the sequon decreased the toxicity of the protein substantially (Fig. 6E), suggesting that CCTX2 exploits the ER protein retention receptor Erd2 for vesicular trafficking. However, it is worth noting that not all CCTX2 homologues share this C-terminal KDEL-like sequon.

Structural data provide further insights into the possible translocation of CCTX2 to the ER. The HSEL sequence is hidden inside the bulk of the C-terminal domain (Fig. 4C), shielded from direct contact with the solvent by an α-helix (residues 589-600). However, the helix carries a putative kexin cleavage site (K596-R597), which is cleaved by the *S. cerevisiae* homologue Kex2p in *in vitro* assays (Fig. S7C). Kexin is a TGN protease and a lower eukaryote orthologue of furin, a peptidase involved in the activation of several toxins (*e.g.*, Shiga toxin) (Garred et al, 1995). The fragments resulting from Kex2p cleavage do not dissociate after proteolysis (Fig. S7C), which can be explained by the highly compact fold of CCTX2. Thus, we hypothesize that proteolytic processing causes a conformational change in the protein, exposing the HSEL sequon to the solvent, allowing recognition by the KDEL receptor and transportation back to the ER.

### CCTX2 effector domain

The nematotoxicity of CCTX2 is dependent on its C-terminal domain of unknown function (Fig. 4B and 4C), thus identifying it as the defense effector domain. Neither BLAST nor structural searches revealed any similarities to proteins with known functions, leaving its biological role unclear. Sequence comparisons among fungal homologues and functional analysis revealed a conserved RxDxQ motif that is essential for the nematotoxicity of CCTX2 (Figs. 1B, S1D and 4C). This finding does, however, not help to draw conclusions regarding the function of the domain as such a motif has, to our knowledge, not been described for any other protein.

The cytotoxic domain adopts an α+β-fold: a large, mixed seven-stranded β-sheet supported by two α-helices constitutes its core folding motif (Fig. 3). Searching the PDB for structural matches did not return any hit, revealing a new protein fold. The β-sheet exhibits a large solvent-exposed surface, which could act as a catalytic cleft, and is decorated by two insertions, mostly composed of α-helices and random coil segments. Site-directed mutagenesis targeting 23 amino acid residues conserved within the CCTX family, identified D687 and Q689 as essential for *C. elegans* toxicity (Fig. 4C and S7). Their placement at the putative catalytic cleft (Fig 3D and 3E) is consistent with a potential role in substrate binding. However, the fold observed in the cryo-EM structure might not be the active form of the effector domain of CCTX2. As discussed in the context of the putative ER retention signal, the proteolytic processing by kexin could trigger a structural rearrangement that considerably alters the domain topology and eventually also the overall fold of CCTX2. Interestingly, an Ala variant targeting residue Asp585, positioned in the high-*B*-factor loop immediately preceding the kexin-cleaved helix, shows a decrease in activity (Fig. S7A and Fig. S8A), suggesting its active role in dynamic changes affecting that region, such as a conformational change.

Finally, several features hint to a role of metals in the DUF domain activity. A putative Zn^2+^-binding site formed by Cys residue 514, 515 and 550 has been detected at the interface between BTF domain 4 and the DUF domain (Fig. 2B). The Cys arrangement suggests a structural, rather than an enzymatic role for the zinc ion (Pace & Weerapana, 2014). The three cysteines are not conserved in CCTX1 and CCTX3; however, they are present in other putative members of the CCTX family (Fig. S1D). More intriguingly, the mutation of Thr739 to an Ala leads to a sharp decrease in toxicity. Thr739 lies at the bottom of a solvent-exposed pocket, placed at the base of the DUF domain (Fig. S8C) and mostly lined by negatively-charged amino acids (Glu786 and aspartates from the DDDxL motif) - an ideal binding spot for divalent cations. Metal-binding has also been found to be essential for the activity of other chimerolectins/toxins, *e*.*g*. MOA and PSL1a (Cordara et al, 2011; Wohlschlager et al, 2011; Cordara et al, 2016; Cordara et al, 2017; Manna et al, 2020).

### Concluding remarks

We provide an in-depth investigation of CCTX2, the holotype of a new family of fungal chimerolectins/AB toxins. While CCTX2 undergoes the same GSL-mediated internalization and retrograde trafficking as many plant and bacterial AB toxins, it reveals a unique domain topology and effector domain. This domain adopts a protein fold not found in any structure deposited in the PDB. Further investigation will be needed to identify the biochemical function of this domain and the detailed molecular mechanism of this protein toxin.

## MATERIALS AND METHODS

### Strains and cultivation conditions

*Escherichia coli* strains DH5α and Turbo (New England Biolabs Inc.) were used for cloning and plasmid amplification, while strains BL21(DE3) and C41(DE3) (Sigma) were used for protein production. *E. coli* strains were cultivated in Luria broth (LB) medium, using well-known protocols described *e.g.* in Sambrook *et al*. (Sambrook & Russell, 2001). *E. coli* strain OP50 was used as food source for *C. elegans* (Stiernagle, 2006). The Bristol isolate N2 was used as the wild-type *C. elegans* strain; *C. elegans* strains were propagated on nematode growth medium (NGM) as previously described (Stiernagle, 2006). All the *C. elegans* strains included in this study were obtained from *Caenorhabditis* Genetics Center (CGC; University of Minnesota) and are listed in Table S8.

### Cloning and site-directed mutagenesis

The full-length coding sequences for CCTX2, CCTX1 and CCTX3 were amplified from *Coprinopsis cinerea* AmutBmut fruiting body (CCTX2, CCTX1) and vegetative mycelium cDNA (CCTX3), using primer pairs CCTX1for_His_NdeI/CCTX1rev_NotI (CCTX1), CCTX2for_NdeI/CCTX2rev_NotI (CCTX2), and CCTX3for_His_NdeI/CCTX3rev_NotI (CCTX3), respectively (Table S9). cDNA synthesis was performed using the Transcriptor Universal cDNA Master kit (Roche) and Phusion DNA polymerase (NEB) as described in Plaza *et al*. (Plaza et al, 2014). The resulting PCR amplicons were cloned into the expression vector pET24b (Novagen) using *Nde*I (Thermo Scientific) and *Not*I (Thermo Scientific) restriction sites. The list of constructs used in this work and their use is reported in Table S7.

#### Constructs for functional studies

The pET24_CCTX2 plasmid served as the template for the construction of N- and C-terminal truncations of the CCTX2 coding region, including the deletion of the C-terminal HSEL sequon, and for the site-directed mutagenesis of specific residues to alanine. Constructs pET24_CCTX2ΔN (lacking residues 2-303) and pET24_CCTX2ΔC (lacking residues 570-787), both with a N-terminal His_8_ tag, were generated using primer pairs CCTX2ΔNfor_His_NdeI/CCTX2rev_NotI and CCTX2for_His_NdeI/CCTX2ΔC_rev_NotI, respectively. The pET24-CCTX2ΔHSEL construct was cloned using primer pair CCTX2for_NdeI/CCTX2_ΔHSEL_rev_NotI (Table S9); the corresponding PCR products were cloned into the pET24b expression vector using *Nde*I and *Not*I restriction sites. Site-directed mutagenesis of pET24-CCTX2 was performed by overlap extension PCR using Phusion DNA polymerase and a combination of CCTX2for_NdeI, CCTX2rev_NotI and mutagenesis primers with the respective single, double and triple mutations. All plasmids were amplified in *E. coli* strain DH5α, purified using mini- and midi-preparation kits (Macherey-Nagel) and verified by Sanger sequencing (Microsynth, Switzerland). Plasmids carrying the correct mutations were then transformed into *E. coli* strain BL21(DE3) for protein production and purification.

#### Constructs for structural studies

A different expression vector was generated for structural studies, carrying CCTX2 fused to a tobacco etch virus (TEV) protease-cleavable His_8_ tag. Starting from the pET24_CCTX2 construct, the *cctx2* gene was subcloned into the pET22b(+) expression vector (Novagen) to generate the pET22b(+)-8H-TEV-CCTX2 construct (Table S7). The cloning was carried out using the NEBuilder HiFi DNA Assembly kit (New England Biolabs), following guidelines from the manufacturer. Cloning primer pairs for the gene (CCTX2_fwd/CCTX2_rev; Table S9) and the vector (pET22_fwd/pET22_rev; Table S9) were designed using the NEBuilder Assembly tool (https://nebuilder.neb.com). PCR was performed using Q5 High-Fidelity DNA Polymerase (New England Biolabs Inc.) following the protocol provided by the manufacturer. Template DNA was degraded using endonuclease *Dpn*I (Thermo Scientific) and assembly of the pET22b(+)-8H-TEV-CCTX2 vector was carried out using the HiFi DNA Assembly Master Mix (New England Biolabs Inc.). Turbo *E. coli* cells (New England Biolabs Inc.) were transformed using the assembly digest and grown o/n at 37°C on LB-agar plates, complemented with 100 μg/mL ampicillin. Single colonies were used to inoculate 5 mL LB media cultures supplemented with 100 μg/mL ampicillin; DNA extraction and purification was performed using the Nucleospin Plasmid kit (Macherey-Nagel). Constructs were verified by Sanger sequencing (Eurofins Genomics GmbH, Germany). Clones carrying plasmids with the correct sequence were amplified by o/n growth at 37°C in 200 mL LB cultures, supplemented with 100 μg/mL ampicillin. DNA extraction and purification for long-term storage was performed using the NucleoBond Xtra Midi kit (Macherey-Nagel).

### Protein production and purification

#### Protein for functional studies

Constructs were transformed into *E. coli* BL21(DE3) for protein production and purification. *E. coli* BL21(DE3) transformants were grown in LB medium to an OD_600_ ≈0.7 at 37°C. Cultures were put on ice for 10 to 15 min before protein production was induced with 0.5 mM IPTG (isopropyl β-D-1-thiogalactopyranoside) for 16 h at 20°C under constant shaking at 120 rpm (Multitron incubator-shaker, Infors HT). All proteins were tested for solubility according to the method described by Künzler *et al*. (Künzler et al, 2010). Bacteria were harvested by centrifugation and the pellets were resuspended in 20 mL of 20 mM HEPES, 500 mM NaCl, pH 7.5 and lysed using a French press (Avantec). Thereafter, the lysates were centrifuged at 16’000 rpm (Sorvall SS34 rotor) for 30 min to remove cell debris. The resulting supernatant, containing the soluble whole protein fraction, was incubated for 1 h at 4°C with Sepharose™ CL 6B (GE Healthcare Life Sciences) beads or Ni-NTA beads (Macherey-Nagel), for untagged and His-tagged proteins, respectively. Proteins were batch eluted from Sepharose™ CL 6B or Ni-NTA beads by gravity flow using lysis buffer supplemented with either 200 mM lactose or 250 mM imidazole, respectively. Proteins were desalted on a PD-10 column (GE Healthcare Life Sciences) and the buffer exchanged for 20 mM HEPES and 150 mM NaCl, pH 7.5. MOA, *Coprinopsis cinerea* galectin 2 (CGL2) and *Coprinopsis cinerea* lectin 2 (CCL2) were expressed and purified as described previously (Butschi et al, 2010; Wohlschlager et al, 2011; Schubert et al, 2012).

#### Protein for structural studies

*E. coli* C41 (DE3) cells were transformed with pET22b(+) following a standard protocol, like the one described *e.g.* in Sambrook *et al*. (Sambrook & Russell, 2001). C41 (DE3) was selected among a panel of several *E. coli* strains, as it minimized CCTX2 cleavage by endogenous proteases. Glycerol stocks were prepared from successful transformants, cultured o/d at 37°C in LB supplemented with 100 µg/mL ampicillin and stored at −80°C. 2 mL precultures were inoculated with multiple colonies, taken from LB-agar plates streaked with glycerol stocks, and grown o/d at 37°C, shaking at 120 rpm (Multitron incubator-shaker, Infors HT). 50 mL LB medium cultures containing 100 µg/mL ampicillin were inoculated with the 2 mL precultures and grown at 37°C, 120 rpm for 16 hours. The 50 mL cultures were then used to inoculate 1 L of LB medium complemented with 100 µg/mL ampicillin to an optical density at 600 nm (OD_600_) of 0.1, and incubated at 37°C, 120 rpm. When OD_600_ reached ≈ 0.6, the culture was cooled down in iced water for 10 to 15 minutes. Expression of the gene was induced with 0.5 mM IPTG for 18-20 hours at 20°C, shaking at 120 rpm (Multitron incubator-shaker, Infors HT).

Cells were harvested at 4,000 x *g*, 4°C for 30 minutes and resuspended in 5 mL of lysis buffer per gram of cell paste. The lysis buffer was prepared to a final concentration of 50 mM HEPES, 200 mM NaCl, 2 mM EDTA, 5 mM DTT; the pH was adjusted to pH 7.5. The buffer was complemented with c*O*mplete EDTA-free protease Inhibitor cocktail (Roche Life Science) to the working concentration recommended by the manufacturer. Cells were lysed using an ultrasound homogenizer (Qsonica Q500) at 20% amplitude for 1 minute (10 second cycles: 3 seconds ‘on’, 7 seconds ‘off’). Cell lysates were clarified at 40,000 x *g*, 4°C for 40 minutes. The supernatants were recovered and directly used for purification.

Galactose-affinity chromatography was carried out at RT on an ÄKTA Start (GE Life Sciences), using Immobilized D-galactose gel (Thermo Scientific) packed into a 5 mL Tricorn 10/50 column (GE Life Sciences). The affinity column was equilibrated with loading buffer (20 mM HEPES, 500 mM NaCl, pH 7.5), loaded with cell lysate and washed with six column volumes (CV) of loading buffer. Bound protein was eluted with ten CV of elution buffer (20 mM HEPES, 500 mM NaCl, 1 M D-galactose, pH 7.5). Fraction content was assessed by SDS-PAGE; fractions containing CCTX2 were pooled and concentrated by ultrafiltration at 3,500 x *g*, 4°C using Vivaspin® 20 Centrifugal Filter Units 10K Molecular Weight Cut Off (MWCO) (VWR). Size exclusion chromatography (SEC) was performed, to ensure sample purity and monodispersity. A Superdex 200 Increase 10/300 GL (Cytiva) gel-filtration column was equilibrated with SEC buffer (50 mM HEPES, 150 mM NaCl, 100 mM Galactose; pH adjusted to 7.5) and loaded with the concentrated sample from ultrafiltration. The content of eluted fractions was assessed by SDS-PAGE; fractions containing pure CCTX2 were pooled and loaded onto a 5 mL HiTrap desalting column (Cytiva), previously equilibrated with dialysis buffer (20 mM HEPES, pH 7.5). The dialysed protein was kept at 4°C until its use for structural studies.

### Biotin, Alexa Fluor®488 and TAMRA protein-labeling

For thin-layer chromatography (TLC), glycan array analysis and *C. elegans in vivo* imaging, proteins were purified as described above (‘Protein purification for functional studies’) and biotinylated with the EZ-Link®-sulfo-NHS-biotin kit (Thermo Scientific, USA). The proteins were labeled following the manufacturer instructions, using a 20-fold molar excess of labeling agent. Protein labeling with Alexa®488 (Thermo Scientific, USA) and carboxytetramethylrhodamine (TAMRA; Thermo Scientific, USA) was performed as described previously (Bleuler-Martínez et al, 2011; Wohlschlager et al, 2011; Stutz et al, 2015).

### Preparation and analysis of glycosphingolipids from *C. elegans*

Total lipids were extracted, and upper phase lipids containing glycosphingolipids (GSLs) were separated by thin-layer chromatography (TLC) according to Barrows *et al*.(Barrows et al, 2006; Wohlschlager et al, 2011). Binding of separated lipids to CCTX2 was assessed by overlaying the TLC plates with biotinylated CCTX2 and HRP-coupled avidin. The biotinylated GSL-binding chimerolectin MOA (Wohlschlager et al, 2011) and the N-terminally truncated version of CCTX2 (CCTX2ΔN) were used as controls. GSLs were extracted from *C. elegans* wild-type N2 strain as well as from the mutant strains *bre-2, bre-3, bre-4* and *bre-5* and separated using TLC, as described by Barrows *et al*. (Barrows et al, 2006). In brief, L4-staged larvae from each strain were collected and washed three times in 10 ml H_2_0. The resulting pellet was flash-frozen in liquid nitrogen and worm pellets were sonicated to break the worm cuticle. 1 volume of homogenized worm pellet was mixed first with methanol and then chloroform to a final ratio of 4:8:3 (chloroform:methanol:water). Samples were incubated for 2 h at 37°C and afterwards centrifuged 5 min at 1400 x *g*. Supernatant was transferred into a new glass vial. 0.147 volumes of water were added to the supernatant. Samples were centrifuged again as above to achieve phase separation. The upper phase (containing the glycolipids) was transferred onto a Sep-Pak cartridge (Millipore) for reversed-phased chromatography. For TLC overlay analysis, 150 nM biotinylated MOA, CCTX2, or CCTX2ΔN were used following the protocol described by Wohlschlager *et al*. (Wohlschlager et al, 2011).

### *C. elegans* toxicity assay in liquid medium with purified protein

*C. elegans* toxicity assays were carried out as previously described (Künzler et al, 2010; Stutz et al, 2015) with the following modifications: Assays were performed in 96-well plates, with approximately 30 L1 staged larvae. Larvae were incubated in 100 µl 1x phosphate-buffered saline (PBS; pH 7.4) containing *E. coli* OP50 OD_600_ = 2 and the indicated protein concentrations for 72 h at 20°C. Larval development into L4 stage or adulthood was scored as full development.

### Microscopy of *C. elegans*

*C. elegans* strains were grown into L4 stage and fed with 150 μg/ml of 5-TAMRA-labeled MOA or CCTX2 or with 500 μg/ml of 5-TAMRA-labeled CCL2 for the indicated time, before washing 3x with 1x PBS to remove excess protein. Worms were immobilized using 10 mM levamisole and mounted on a 3% w/v agarose pad. Fluorescence images were taken using a Zeiss LSM 780 upright confocal microscope. Image acquisition was performed using the ZEN system imaging software (Zeiss, Germany). Image analysis was performed using the software Fiji (Schindelin et al, 2012), a distribution of the open-source imaging software ImageJ (Schneider et al, 2012).

### Kex2p proteolysis assay

100 μg of CCTX2 were incubated at 37°C overnight (o/n) with 10 μg recombinant Kex2p from *Saccharomyces cerevisiae* (PeproTech). Samples from the reactions were collected, diluted 1:10 and separated by SDS-PAGE. Gel bands were excised and identified by tryptic digest and peptidomics (Functional Genomics Center Zürich). The remainder of the reactions was incubated with Sepharose^TM^ CL 6B or Ni-NTA beads for 1 h at RT on a rotating wheel. Thereafter, the slurry was loaded onto Mobicol spin columns (MoBiTec, Germany) and the flow-through was collected. The beads were washed with 50 column volumes of 20 mM HEPES, 150 mM NaCl, pH 7.5 and boiled in Laemmli buffer. The eluates were subjected to an immunoblot analysis using a polyclonal rabbit antiserum raised against recombinant CCTX2 (Seramun Diagnostica; Fig. S7C).

### Sequence analysis, phylogeny and structure prediction

Domain predictions were performed using the SMART software (Simple Modular Architecture Research Tool, biobyte solutions GmbH, Germany; http://smart.embl-heidelberg.de). CLUSTALW amino acid sequence alignments were performed using the Pole Bio-Informatique Lyonnais (PBIL)-based software NPS@CLUSTALW (Combet et al, 2000) (Network Protein Sequence @nalysis, Lyon, France; https://npsa-prabi.ibcp.fr/NPSA/npsa_clustalw.html). The multiple sequence alignment of the three *C. cinerea* paralogues CCTX1, CCTX2 and CCTX3, as well as their fungal full-length homologues, was done using the MUltiple Sequence Comparison by Log-Expectation (MUSCLE) algorithm (Edgar, 2004). The alignment was visualized in Unipro UGENE software (Okonechnikov et al, 2012) and manually trimmed (Fig. S1D). The phylogenetic tree was constructed using the RAxML-ng algorithm (Kozlov et al, 2019) which implements maximum-likelihood (ML) optimality criterion, setting 500 bootstrap replicates under a Jones-Taylor-Thornton matrix. The phylogenetic tree with the highest bootstrap support was visualized (Fig. 1B) using ITOL (Letunic & Bork, 2024). *In silico* structure prediction for CCTX2 was initially carried out using the Robetta server (Kim et al, 2004) (https://robetta.bakerlab.org), based on the Rosetta simulation framework (Baek et al, 2021), applying the ‘comparative modeling’ protocol and automatic template selection (Song et al, 2013b). Later on, a model was generated using AlphaFold 3 (Abramson et al, 2024). Homology models of CCTX1 and CCTX3, used for domain boundary prediction, were generated using the Robetta server (Kim et al, 2004) (https://robetta.bakerlab.org), applying the ‘comparative modeling’ protocol and uploading the cryo-EM structure of CCTX2 as a template.

### Cryo-EM grid preparation and data collection

Cryo-EM grids were prepared at the SciLife lab cryo-EM Swedish National facility, Umeå node (Sweden) according to standard procedures described in *e.g.* Passmore and Russo (Passmore & Russo, 2016). Quantifoil R2/1 300 mesh carbon grids (Quantifoil Micro Tools, Germany) were glow-discharged at 15 μA for 30 s with a Pelco easiGlow glow-discharge cleaning system (Ted Pella, Inc.). 4 μL of 1 mg/mL CCTX2 were applied to the grids using a Vitrobot Mark IV (Thermo Scientific) equilibrated at 4°C, 100% humidity. 7,500 movies were collected on a FEI Titan Krios (Thermo Scientific) operating at 300 kV and equipped with a Gatan K2 camera, at a nominal magnification of 215,000 x and with a physical pixel size of 0.63 Å. The dose rate was set to 4.7 e^−^/px/s, and the total exposure time was 5 s, resulting in a total dose of 59.5 e^−^/Å^2^. Nominal defocus range was −1.5 μm to −3.0 μm in 0.3 μm steps. Data collection statistics are reported in Table S2.

### Image processing and model building

The data set was processed using *cryoSPARC* 3 (Punjani et al, 2017) as described in Fig. S4. In brief, patch motion correction and contrast transfer function (CTF) estimation was performed. After manual particle picking, homogenous refinement was carried out to generate a volume for template-based particle picking, generating a stack of 1.8 million particles. After three rounds of 2D classification, 641,000 particles remained with a final extraction at a box size of 480 pixels and cropping to 256 pixels. These particles were used for *ab initio* reconstruction, where the best class was chosen for heterogeneous refinement. 204,000 particles were selected from two rounds of heterogeneous refinement. The volume was subjected to local refinement and sharpening to increase the quality of the map, which extended to 3.2 Å resolution.

Predictive models were generated using the Robetta server (Kim et al, 2004) for all the CCTX2 domains, using either template-based (BTF domains 1 to 4) or with *ab initio* (C-terminal domain) protocols. Directly fitting the models into the reconstructed volume proved impossible, due to the strong structural homology among the BTF domains (plus the presence of a pseudo-threefold axis) and the poor quality of the C-terminal domain prediction; automated protein chain building was not successful either. However, a polyalanine trace, manually built using *Coot* (Casañal et al, 2020), a component of the *CCP4* software suite (Agirre et al, 2023), revealed the correct domain topology. The first three BTF domains were easily docked onto their respective polyAla trace, whereas the fourth domain had to be manually aligned in PyMOL (Schrödinger, Inc.), only relying on the backbone segments built into the discontinuous Coulomb potential density (Fig. S3A). Poorly predicted segments were removed from the model of the C-terminal domain, keeping only a core region (residues 619-740; Figs. S3B and S3C). The CCTX2 model was completed using *Coot*, by accurately fitting the models into the cryo-EM volume and building missing residues; an *in silico* model generated using AlphaFold 3 (Abramson et al, 2024) was used to improve areas with poor density fit. Refinement was carried out by alternating cycles of real-space building using *Coot* (Casañal et al, 2020) and real-space refinement with *phenix.refine*, a component of the *Phenix* software suite for structural biology (Liebschner et al, 2019). The final model includes the entire CCTX2 chain (residues 1-787); the C-terminal amino acid residues from the His_8_-TEV tag were poorly defined and were not modeled. Similarly, galactose molecules at the putative sugar-binding sites of the first (subdomains β and γ) and second (subdomain α) BTF domains were not modeled due to poor fit density. Refinement statistics are reported in Table S2: although the model displays a remarkably high number of Ramachandran outliers (8, or about the 1%, evaluated with RAMPAGE (Lovell et al, 2003)), their geometry is generally close to the allowed regions of the plot. All structural illustrations were prepared using PyMOL (Schrödinger, Inc.).

## ACKNOWLEDGEMENTS

We would like to thank Ken Teter for comments on the manuscript. Cryo-EM experiments were performed at SciLife lab (Umeå, Sweden), many thanks to Michael Hall for collecting the cryo-EM data. The structural biology work was carried out at the UiO Structural Biology core facilities, which are part of the Norwegian Macromolecular Crystallography Consortium (NORCRYST) and received funding from the Norwegian INFRASTRUKTUR-program (project no. 245828) as well as from UiO (core facility funds). The genetics, biochemistry and cell biology work was carried out at ETH Zürich and funded by the Swiss National Science Foundation (Grants no. 31003A-130671 and 310030-212435). The PhD position of F.K. was financed by UiO.

## AUTHOR CONTRIBUTIONS

M.K. conceived the study, and G.C., S.S.S. and U.K. additionally contributed to designing the experiments. S.S.S., K.B. and D.P. produced and purified the recombinant CCTX1-3 variants and performed the toxicity and cell biology assays, B.S. prepared the sequence alignments and phylogenetic tree, under the supervision of M.K.. C.H.K. and F.K. produced and purified recombinant CCTX2 for structural biology studies, under the supervision of G.C. and U.K.. F.K. and G.C. analyzed the cryo-EM data, in collaboration with and supervised by A.A.A., A.B., J.L.K. and T.B.. The manuscript was initially written by S.S.S. and M.K., substantially revised by G.C., F.K. and U.K., and finalized by all authors.

## ABBREVIATIONS LIST

BSA: bovine serum albumin
BTF: β-trefoil fold
CCL2: *Coprinopsis cinerea* lectin 2
CCTX1: *Coprinopsis cinerea* toxin 1
CCTX2: *Coprinopsis cinerea* toxin 2
CCTX3: *Coprinopsis cinerea* toxin 3
CGL1: *Coprinopsis cinerea* galectin 1
CGL2: *Coprinopsis cinerea* galectin 2
cryo-EM: cryogenic electron microscopy
CV: column volumes
DUF: Domain of Unknown Function
EDTA: ethylenediaminetetraacetic acid
ER: endoplasmic reticulum
GSL: glycosphingolipid
HEPES: 4-(2-hydroxyethyl)piperazine-1-ethane-sulfonic acid
IPTG: isopropyl β-D-1-thiogalactopyranoside
Kex-2: kexin 2 from *Saccharomyces cerevisiae*
LB: lysogeny broth
MANS: α-Mannosidase II
MOA: *Marasmius oreades* agglutinin
MS: mass spectrometry
MWCO: molecular weight cut-off
NAD: nicotinamide adenine dinucleotide
NGM: nematode growth medium
o/d: over day
o/n: overnight
OD: optical density
PARP: poly(ADP-ribose) polymerase
PBS: phosphate-buffered saline
PDB: Protein Data Bank
rpm: rotation/round per minute
TAMRA: tetramethylrhodamine
TLC: thin-layer chromatography
TGN: trans-Golgi network

