## Supplementary Figures and Tables for "Structure and function of a fungal AB toxin-like chimerolectin involved in anti-nematode defense"

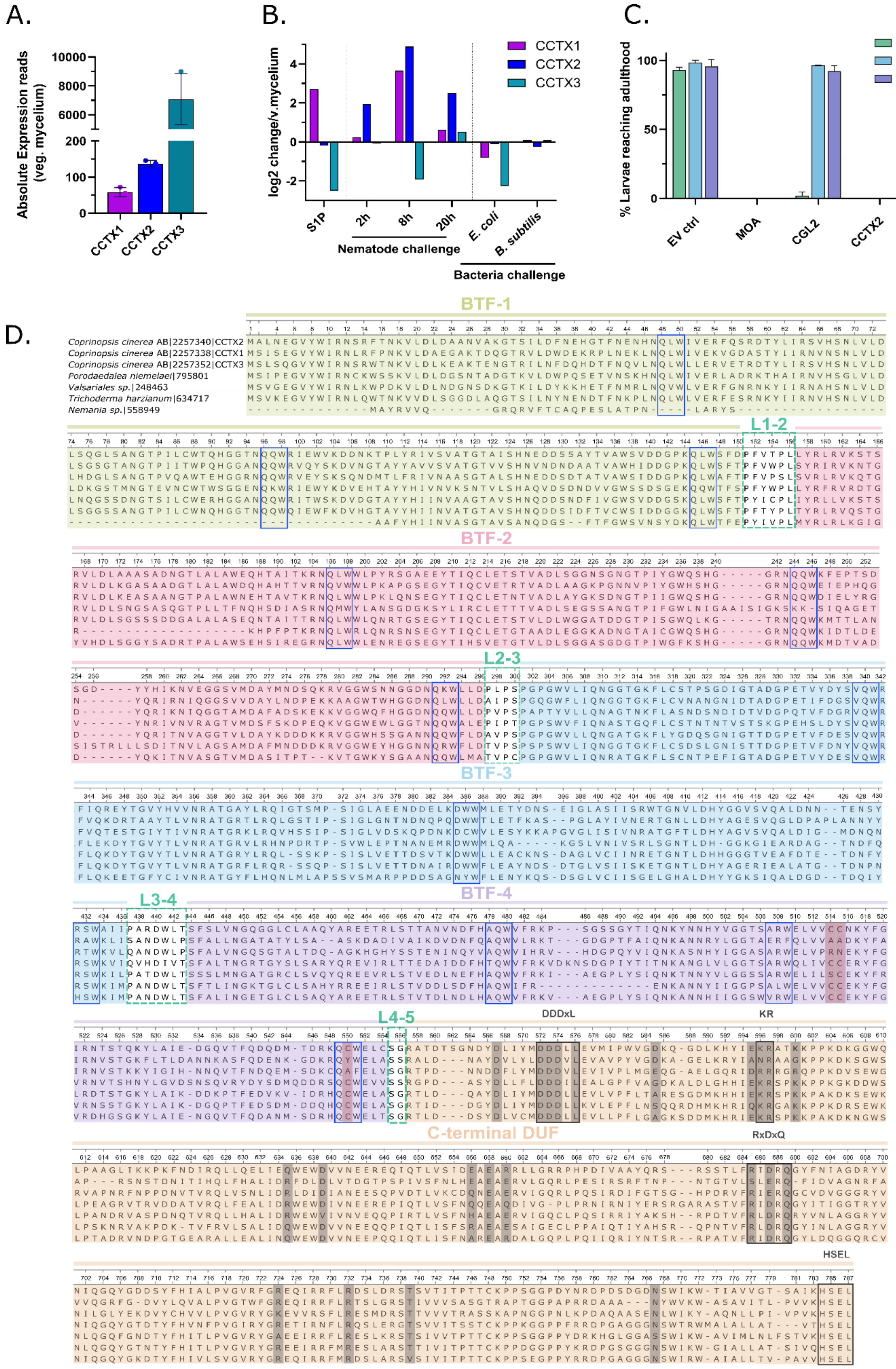

**Figure S1. Differential expression, toxicity and sequence alignment of CCTX2 paralogues**. **A**. Absolute mRNA levels for CCTX1, CCTX2 and CCTX3 in *C. cinerea* vegetative mycelium. Data points with error bars are means of N = 3 biological replicates with standard deviation (SD). Data was taken from Muraguchi et al. (Muraguchi et al, 2015). **B.** Differential mRNA expression analysis of CCTX1, CCTX2 and CCTX3 in *C. cinerea* vegetative mycelium vs S1P primordia and vegetative mycelium challenged by fungivorous nematode *A. avenae*, and two bacterial species, *E. coli* and *B. subtilis*. Data are replotted from ´ (Muraguchi et al, 2015; Tayyrov et al, 2018; Kombrink et al, 2019). **C**. Differential toxicity of the chimerolectins CCTX2, MOA and the lectin CGL2 against three bacterivorous *Caenorhabditis* species N2 (*elegans*), AF16 (*briggsae*) and JU1373 (*tropicalis*). Here, genes were overexpressed in *E. coli* and the recombinant bacteria fed to the nematodes. **D.** Sequence alignment of members of the monophyletic clade of fungal full-length homologues including the three paralogues from *C. cinerea* (see complete results of BLAST search in Fig. 1A). The alignment shows that the overall chimeric architecture of *C. cinerea* proteins, consisting of four lectin domains and a C-terminal domain of unknown function, as well as a DDDxL motif, a putative kexin-cleavage site (KR), a RxDxQ motif and a putative ER-retention signal (HSEL) are conserved among these homologues. Putative zinc binding sites, which are conserved among homologues, are indicated by red shading, while conserved residues that were subjected to single substitutions by alanine in CCTX2 for functional analysis (alanine scanning), are highlighted in grey.

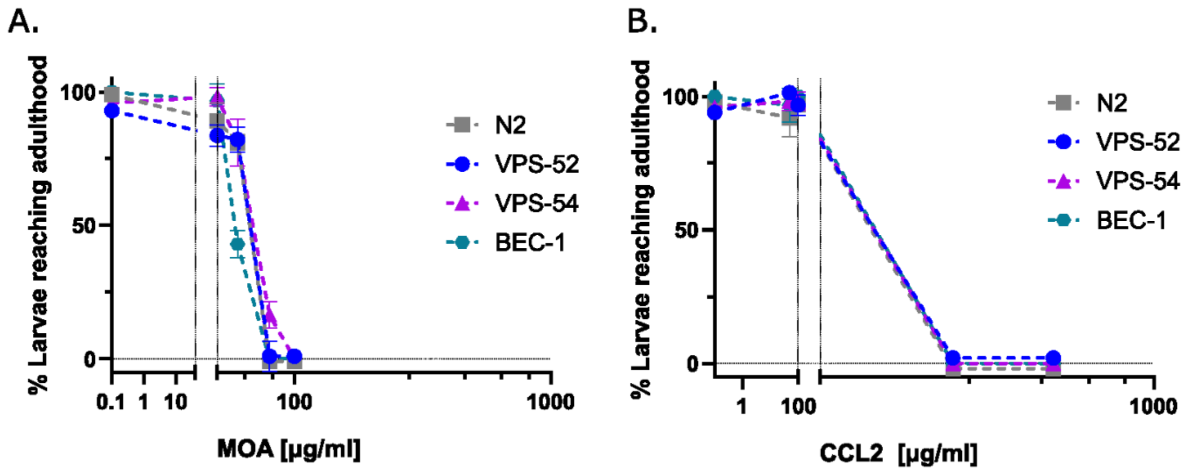


**Figure S2. Toxicity profile of MOA and CCL2 in *C. elegans* endocytosis mutants.** Toxicity assay with CCTX2 and different *C. elegans* retrograde trafficking mutants VPS-52 (VC625) and VPS-54 (VC985) bear mutations in the GARP complex subunits VPS-52 and VPS-54, respectively. BEC-1 is required for retromer localization to endosomes. VPS-36 (VC947) exhibits a mutation in the ESCRT II subunit. **A.** Liquid toxicity assay, performed with *C. elegans* retrograde transport mutants using different concentrations of the purified GSL binding lectin MOA. **B.** The same assay was performed with the N-glycan binding lectin CCL2 using 50, 100, 250 and 500 µg/ml of protein. Data points with error bars indicate means of N = 4 biological replicates with standard error of the mean (SEM). Overlapping data points were nudged by +/- 2 data units in Y for better readability.

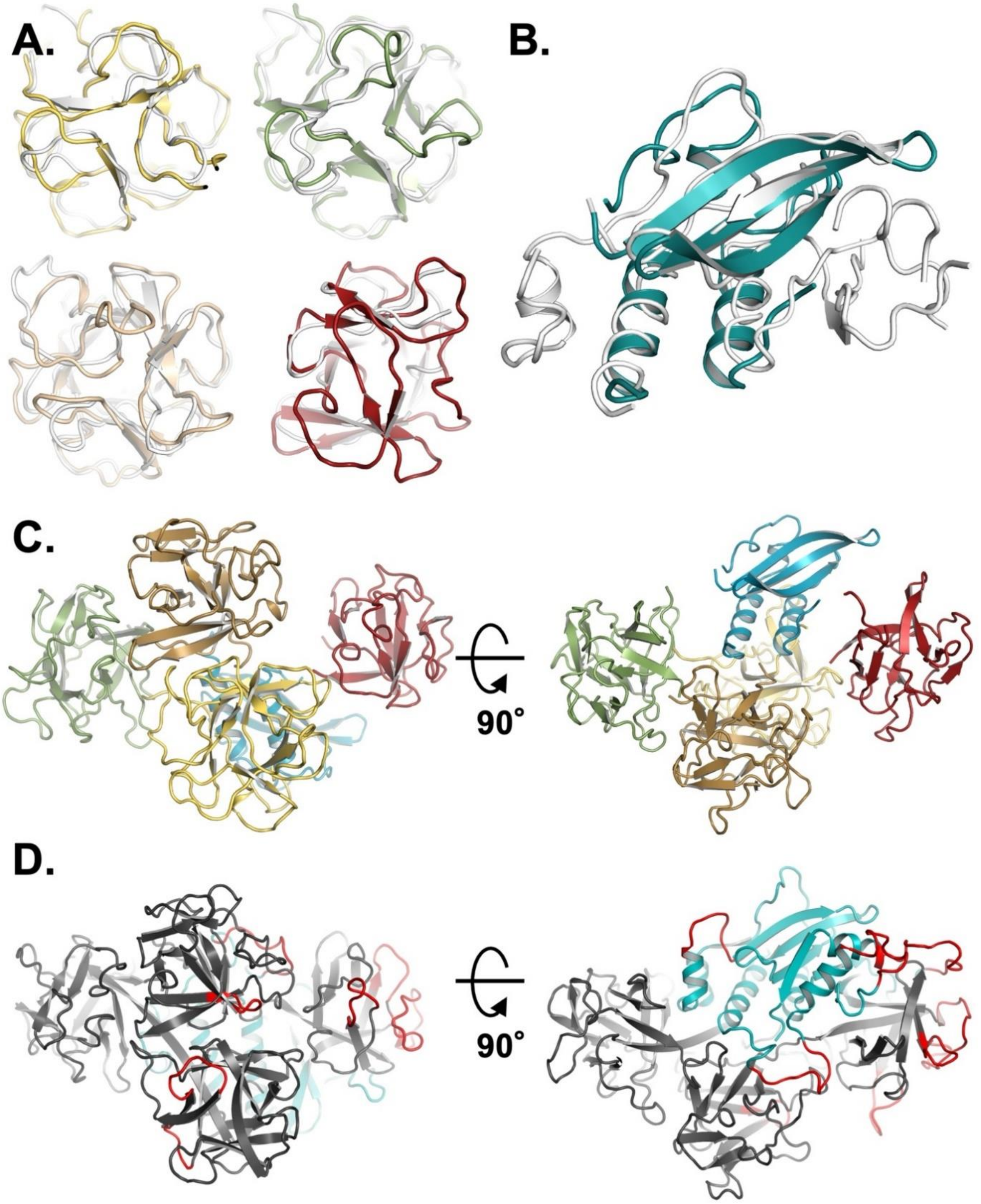


**Figure S3. Model building and comparison of BTF domains. A.** Cartoon representation of the structural alignments of the manually-built polyalanine trace (white) and the template-based homology models generated with Robetta (Kim et al, 2004) for domain 1 (yellow), domain 2 (green), domain 3 (wheat) and domain 4 (red). **B.** Cartoon representation of the structural alignment between the manually built polyalanine trace (white) and the core portion (amino acids 619-740) of the model for the C-terminal domain (deep teal). The model was generated using the Robetta server, applying the *ab initio* protocol and removing all the poorly predicted regions. **C.** Cartoon representation of CCTX2 starting model, after docking all the domain models onto the polyalanine trail. **D.** Final model of CCTX2, with the BTF domains colored in black and the C-terminal DUF domain colored in teal. Highlighted in red are the parts of the model built into partially discontinuous density; boundaries for each region are reported in Table S3.

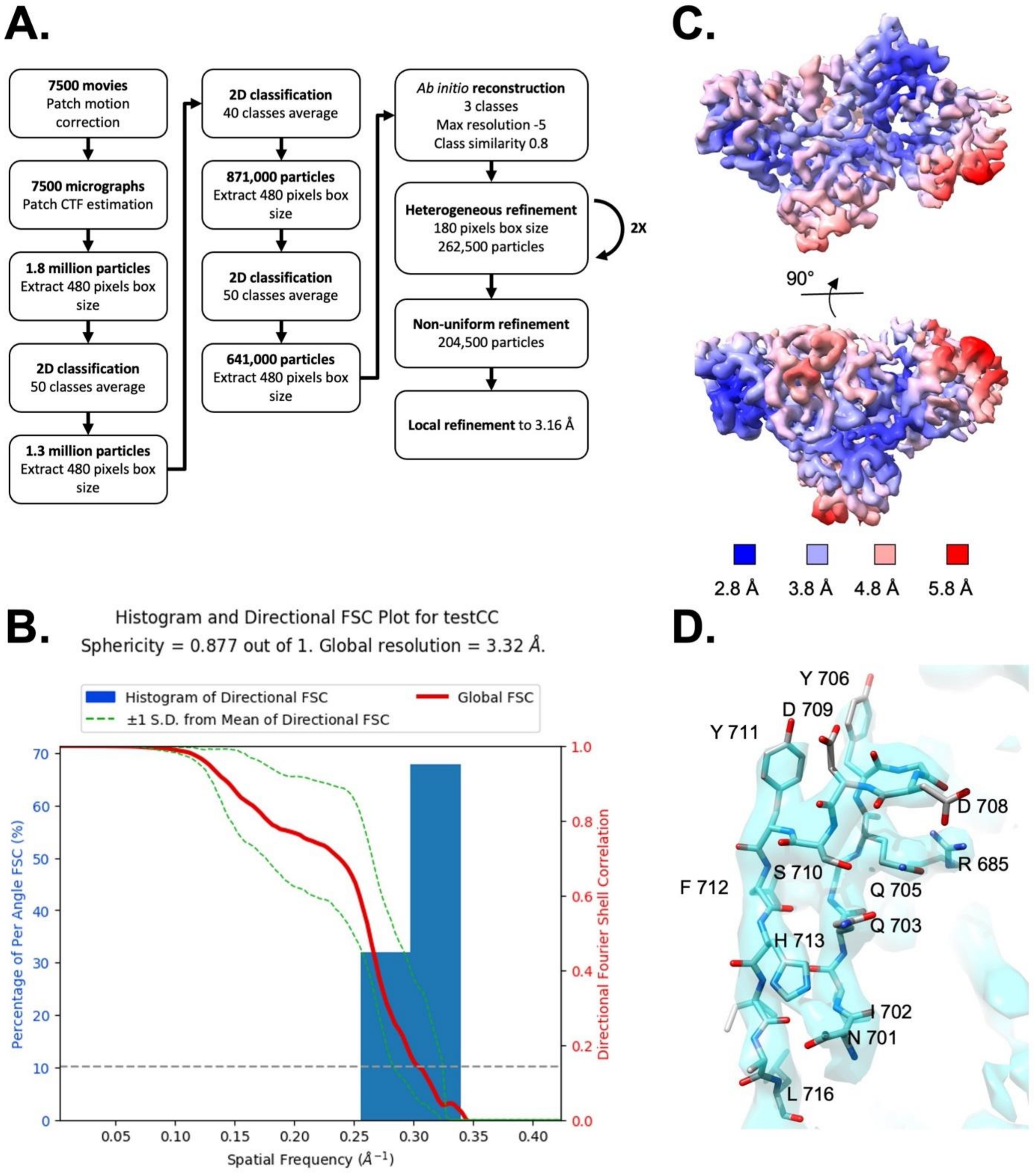


**Figure S4. CCTX2 cryo-EM data processing. A.** Processing workflow in CryoSPARC, leading to the reconstructed cryo-EM volume used to build the CCTX2 model. **B.** Histogram and directional FSC plot for testCC. **C.** Local resolution map. **D.** Example of the quality of the reconstructed coulomb potential map at a loop in the BTF domain 1.

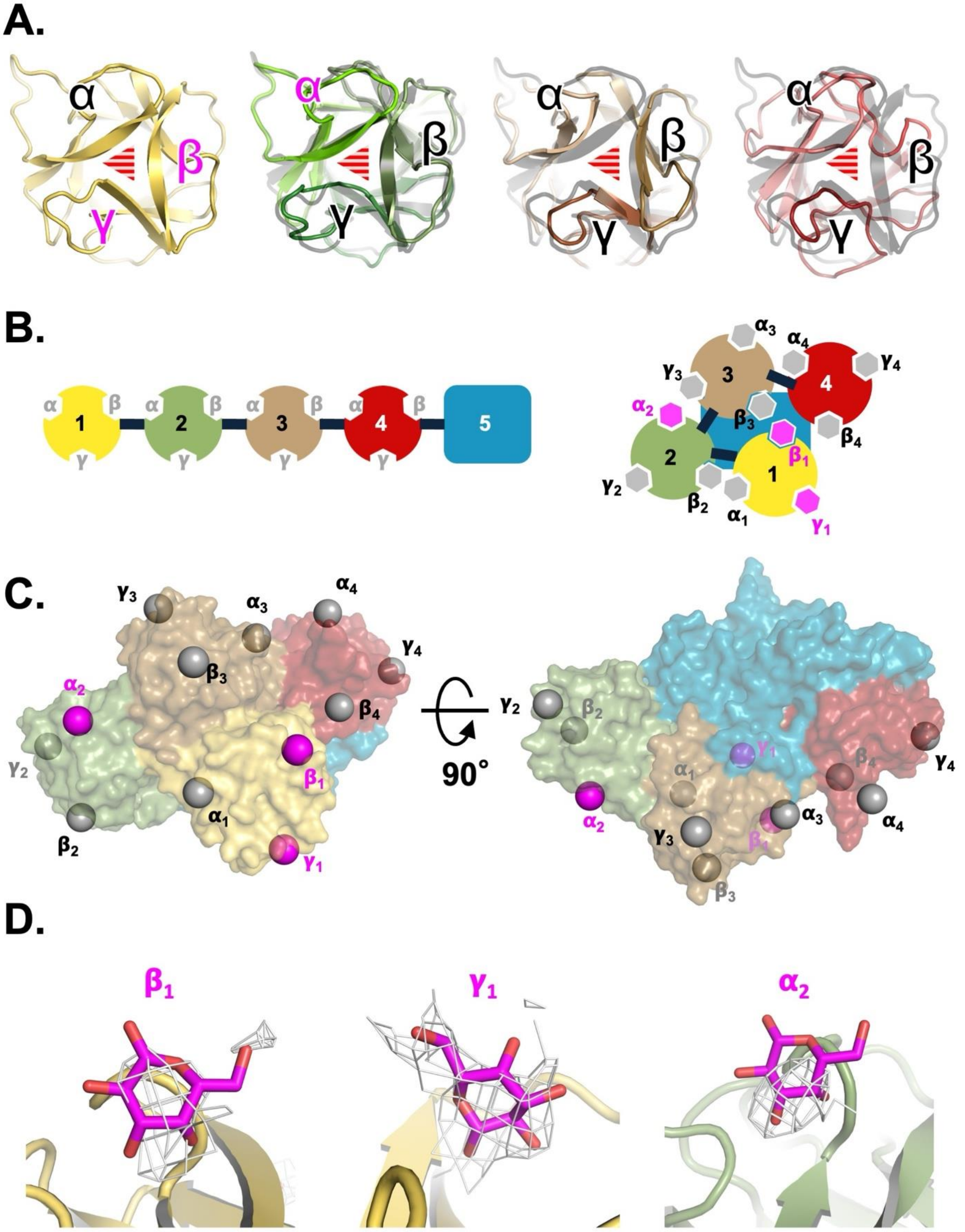


**Figure S5. Beta-trefoil fold (BTF) domains of CCTX2. A.** Cartoon representation of BTF domains 1 (yellow), 2 (green), 3 (wheat) and 4 (dark red). α, β and γ subdomains are marked on each BTF domain; subdomains showing ligand density are marked in magenta. Coordinates of BTF domains 2 to 4 are shown superposed to those of BTF domain 1 (grey) for comparison. The pseudo-threefold symmetry axis associated with the BTF fold is represented by a striped triangle at the center of each domain. **B.** Schematic representation of CCTX2, with the positions of α, β and γ subdomains marked on the BTF domains. The right panel shows a schematic drawing of CCTX2, with the positions of all possible sugar-binding sites marked on the solventexposed surface of the BTF domain cradle. Sugar-binding sites showing Coulomb potential density are marked in magenta. **C.** Surface representation of CCTX2. Putative carbohydrate-binding sites are marked with a grey-colored sphere, while those displaying experimental density are marked with a magenta-colored sphere. **D.** Unmodelled patch of density at the putative sugar binding sites of BTF subdomain $\beta_1$, $\gamma_1$ (left and central) and $\alpha_2$ (right).

A.

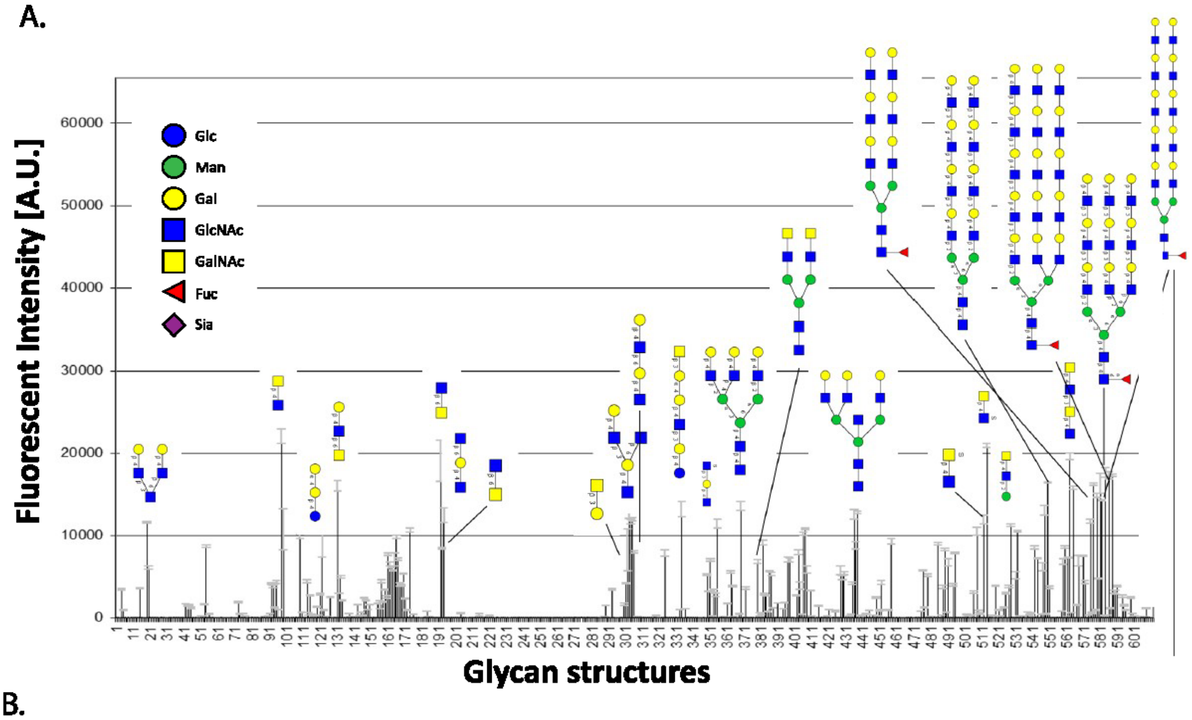


B.

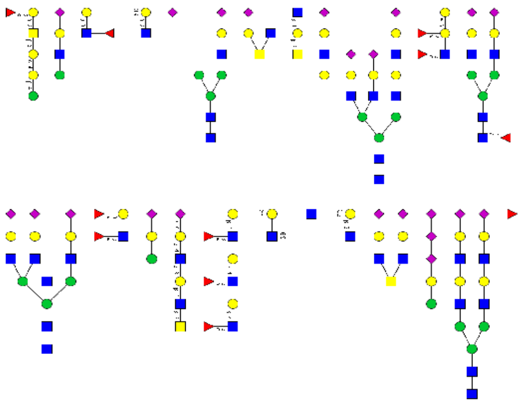

**Figure S6. Glycan array analysis of CCTX2 and the N-terminally truncated CCTX2ΔN**. Purified CCTX2 protein was fluorescently labeled with Alexa Fluor 488 (Invitrogen). The labeled protein was used, at various concentrations, to screen the mammalian glycan array (version 5.1) of Core H of the Consortium for Functional Glycomics (CFG). The data of this analysis is provided as supplementary data files. **A.** Histogram displaying the binding of CCTX2 at a concentration of 20 µg/ml to the glycan array. Glycans with the highest affinity, containing terminal LacNAc and LacDiNAc, are represented as symbols according to (Varki et al, 2015). **B.** Exemplary glycan structures from glycan array screen that contain LacNAc and LacDiNAc motives but are not bound by CCTX2. In these cases, the motives are decorated by additional monosaccharides.

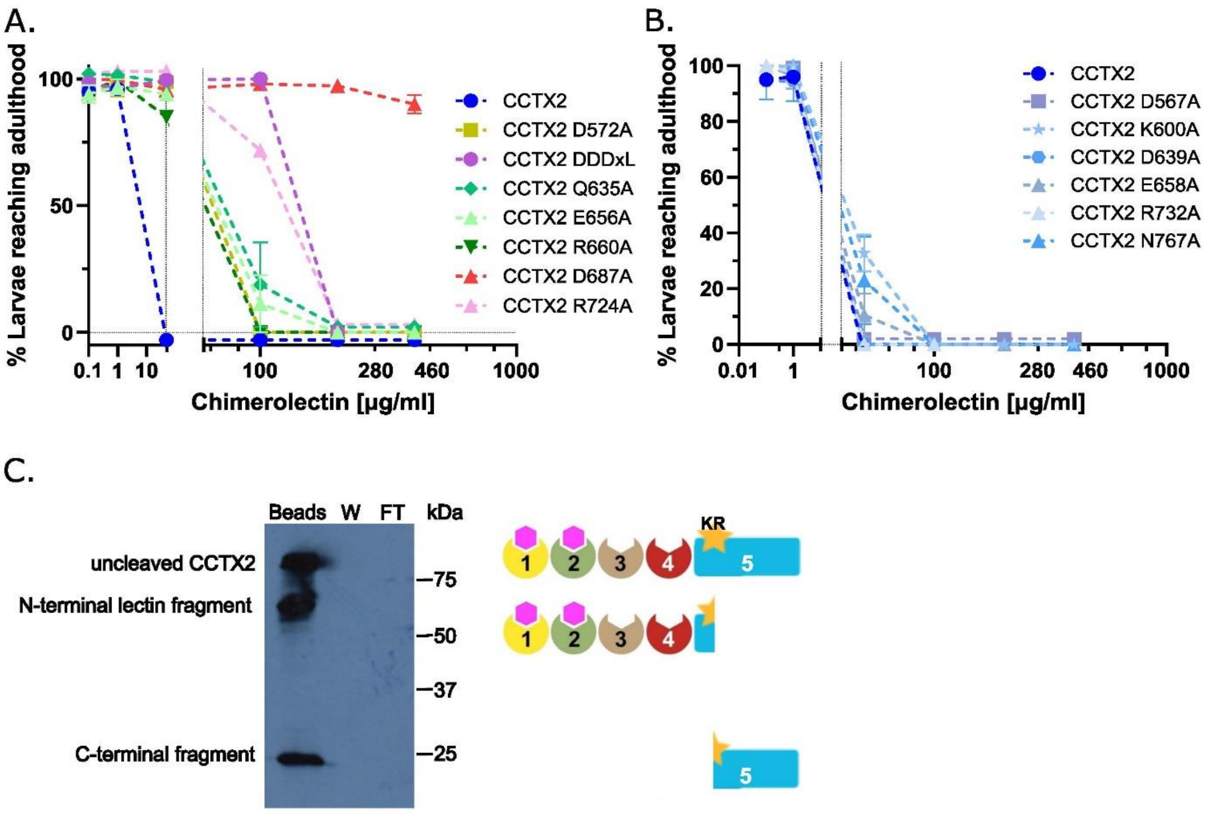


**Figure S7. Alanine scanning of conserved residues within the C-terminal domain of unknown function (DUF). A.** Nematoxicity assay with CCTX2 variants carrying one of seven single residue substitutions that conferred a moderate resistance to the toxin (>100 to <400 µg/ml). WT CCTX2 (dark blue) and D687A, which is part of the 'RxDxQ' motif (red) were included as references. **B.** Nematoxicity assay with CCTX2 variants carrying one of six single residue substitutions that did not reduce toxicity compared to wildtype CCTX2. Data points with error bars indicate means of N = 4 biological replicates with standard error of the mean (SEM). Overlapping data points were nudged by +/- 2 data units in Y for better readability. **C**. *In vitro* cleavage assay of CCTX2 with recombinant *Saccharomyces cerevisiae* Kex2p protease. Kex2p-digested CCTX2 was subjected to a pulldown experiment using Sepharose beads. An immunoblot, using a custom-made anti-CCTX antiserum, of the various fractions run out on SDS-PAGE, shows that all the protein was found in the pulled down fraction ('Beads' lane), including the Kex2p-generated C-terminal fragment, which should be devoid of any lectin domains (BTF1-4). Conversely, no CCTX2 was detected in the wash ('W' lane) or flow-through ('FT' lane) fractions. Two CCTX2 proteolytic fragments (23 and 67 kDa), generated by the Kex2p digestion, are labeled on the left and shown as cartoons on the right side of the gel picture.

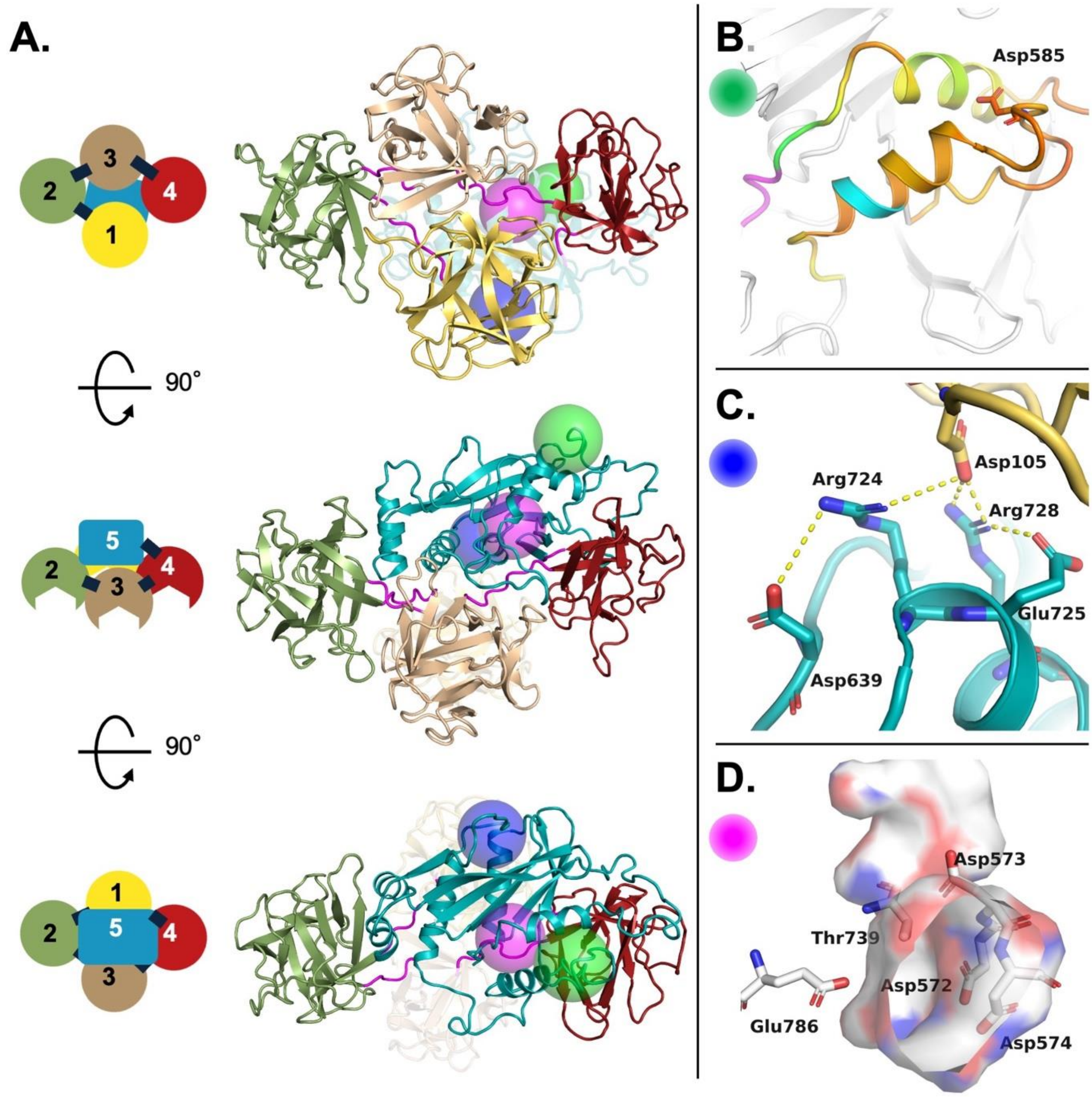


**Figure S8. Structural details of functionally-relevant Ala mutations. A.** Cartoon representation of the CCTX2 structure in three different orientations, which are reprised by a schematic view on the left column; a green sphere marks the position of Asp585, a blue sphere that of Arg724 and a magenta sphere that of Thr739. **B.** Position of Asp585 on the loop preceding helix-1; helix-1 and helix-6 are colored according to their Cα *B*-factors, with the kexin cleavage site marked in cyan and the HSEL sequon in magenta. **C.** Interaction between Arg724 and the surrounding residues, suggesting its possible role in stabilizing the CCTX2 fold by mediating the interaction between the DUF domain (teal) and BTF domain 1 (yellow). **C.** Thr739 lying at the bottom of a pocket, lined by the negatively-charged Glu786 and aspartate residues from the DDDxL motif (Asp572 Asp573 and Asp574).

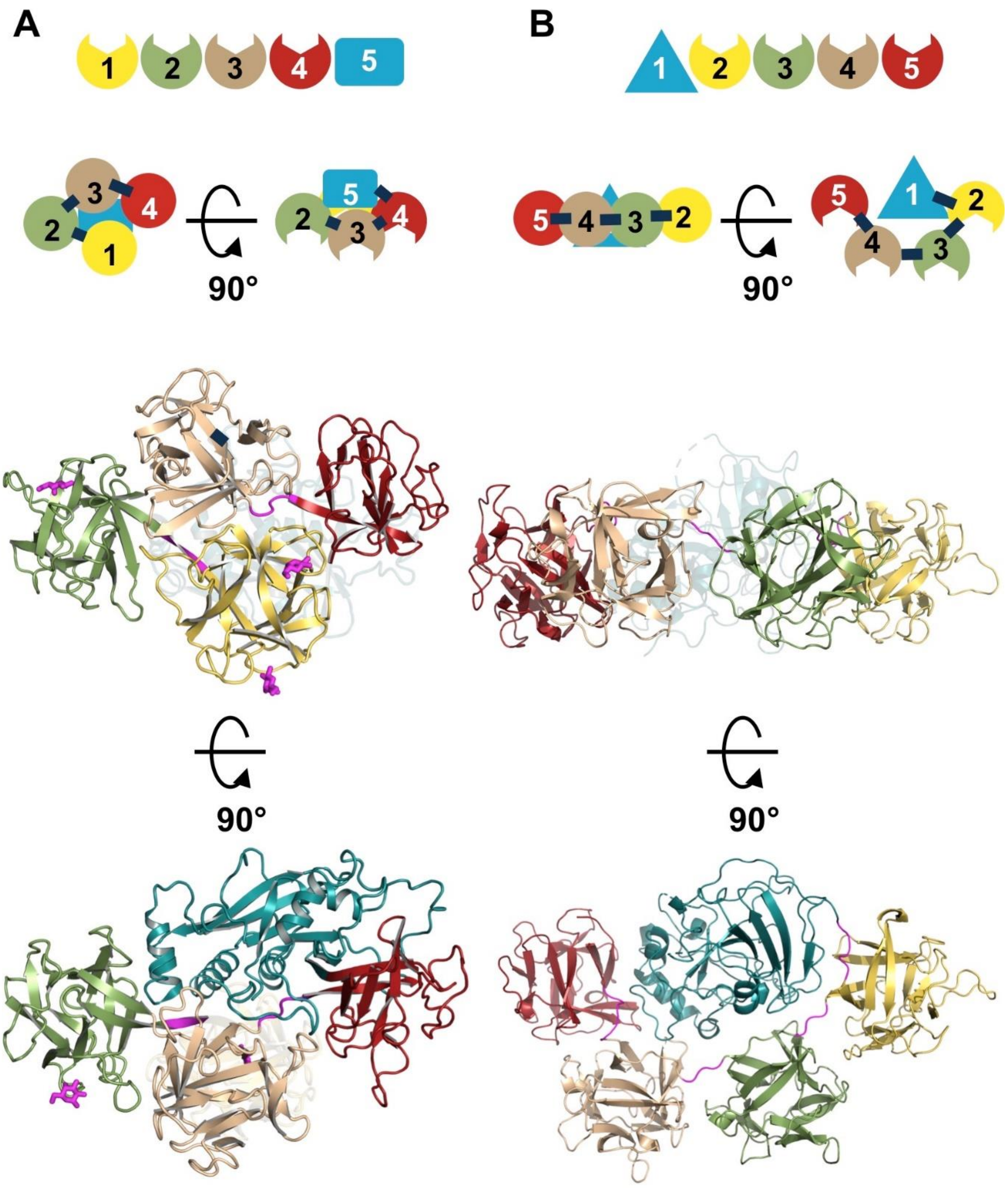


**Figure S9. Comparison of CCTX2 with the mosquitocidal holotoxin from *Pieris rapae* (MTX)**. **A.** Schematic diagram of the domains of CCTX2 and their topology in the protein tertiary structure (PDB ID: 9S11, this work). In magenta: interdomain linkers and D-galactose likely retained from the purification process, modeled into weak density. **B.** Schematic diagram of the domains of MTX and their topology in the protein tertiary structure (PDB ID: 2VSA (Treiber et al, 2008)). In magenta: interdomain linkers.

### Supplementary Tables

**Table S1. List of full-length CCTX2 homologs in the fungal kingdom**

| Name and number of homologues on the tree | JGI Genome Assembly ID | JGI Protein ID |
|---|---|---|
| *Coprinopsis cinerea* AB CCTX1 | Coprinopsis cinerea AmutBmut v2.0 | 2257338 |
| *Coprinopsis cinerea* AB CCTX2 | Coprinopsis cinerea AmutBmut v2.0 | 2257340 |
| *Coprinopsis cinerea* AB CCTX3 | Coprinopsis cinerea AmutBmut v2.0 | 2257352 |
| *Nemania sp.* | Nemania sp. FL0916 v1.0 | 558949 |
| *Valsariales sp.* | Valsariales sp. FL0756 v1.0 | 248463 |
| *Trichoderma harzianum* | Trichoderma harzianum T22 v1.0 | 634717 |
| *Pterula gracilis* | Pterula gracilis CBS309.79 v1.0 | 384083 |
| *Stereum hirsutum* | Stereum hirsutum FP-91666 SS1 v1.0 | 116459 |
| *Choiromyces venosus* 1 | Choiromyces venosus 120613-1 v1.0 | 1801069 |
| *Choiromyces venosus* 2 | Choiromyces venosus 120613-1 v1.0 | 1809985 |
| *Earliella scabrosa* | Earliella scabrosa CIRM-BRFM 1817 v1.0 | 854234 |
| *Ceriporiopsis subvermispora* B 1 | Ceriporiopsis (Gelatoporia) subvermispora B | 125532 |
| *Ceriporiopsis subvermispora* B 2 | Ceriporiopsis (Gelatoporia) subvermispora B | 97606 |
| *Ceriporiopsis subvermispora* B 3 | Ceriporiopsis (Gelatoporia) subvermispora B | 117225 |
| *Ceriporiopsis subvermispora* B 4 | Ceriporiopsis (Gelatoporia) subvermispora B | 117217 |
| *Sistotrema sernanderi* | Sistotrema sernanderi OMC 1753 v1.0 | 84892 |
| *Mycena pura* | Mycena pura 9144 v1.0 | 592943 |
| *Phlebia subcretacea* 1 | Phlebia subcretacea OMC74 v1.0 | 1018667 |
| *Phlebia subcretacea* 2 | Phlebia subcretacea OMC74 v1.0 | 2053875 |
| *Pisolithus orientalis* | Pisolithus orientalis OTSU v2.0 | 900110 |
| *Pisolithus sp.* 1 | Pisolithus sp. B1 v1.0 | 617230 |
| *Pisolithus sp.* 2 | Pisolithus sp. B1 v1.0 | 1098360 |
| *Pisolithus albus* 1 | Pisolithus albus SI12 v1.0 | 2367539 |
| *Pisolithus albus* 2 | Pisolithus albus SI12 v1.0 | 2358149 |
| *Pisolithus microcarpus* | Pisolithus microcarpus 441 v2.0 | 17071 |
| *Pisolithus croceorrhizus* ssp. 2 1 | Pisolithus croceorrhizus ssp. 2 72 v1.0 | 2234632 |
| *Pisolithus croceorrhizus* ssp. A 1 | Pisolithus croceorrhizus subspA 74A v1.0 | 1628709 |
| *Pisolithus croceorrhizus* ssp. A 2 | Pisolithus croceorrhizus subspA 74A v1.1 | 1583998 |
| *Pisolithus croceorrhizus* ssp. 2 2 | Pisolithus croceorrhizus ssp. 2 72 v1.0 | 1889515 |
| *Pisolithus thermaeus* | Pisolithus thermaeus 11 v1.0 | 1506882 |
| *Porodaedalea niemelaei* | Porodaedalea niemelaei PN71-100-IP13 v1.0 | 795801 |

### Table S2 – Cryo-EM data collection and model refinement statistics

| | | |
|---|---|---|
| **Data collection and processing** | | |
| **EMDB ID** | EMD-54430 | |
| **Microscope** | FEI Titan Krios | |
| **Detector** | Gatan K2 | |
| **Voltage (kV)** | 300 | |
| **Magnification (nominal)** | 215,000 | |
| **Pixel size (Å)** | 0.63 | |
| **Electron exposure ($e^-$/Å$^2$)** | 59.5 | |
| **Defocus range (µm)** | -1.5 to -3.0 | |
| **Micrographs collected** | 7,500 | |
| **Micrographs used** | 7,498 | |
| **Particles extracted** | 641,000 / 204,500 | |
| **Simmetry imposed** | n/a (C1) | |
| **Map sharpening** | Global *B*-factor sharpening | |
| **Map resolution (Å)** | 3.16 | |
| **FSC threshold** | 0.143 | |
| **Refinement** | | |
| **PDB ID** | 9S11 | |
| **Initial model used** | Robetta-generated (Kim et al, 2004), template-based and *ab initio* AlphaFold 3 (Abramson et al, 2024) | |
| **Box** | | |
| Pixels | 256, 256, 256 | |
| Angles (°) | 90.00, 90.00, 90.00 | |
| **Pixel size (Å) (map)** | 1.18 | |
| **Resolution estimates (Å)** | **Masked** | **Unmasked** |
| $d_{FSC}$ (half maps; 0.143) | 3.3 | 3.5 |
| $d_{99}$ (full/half$_1$/half$_2$) | 4.3/3.6/3.6 | 4.2/2.7/2.7 |
| d model | 3.9 | 3.9 |
| $d_{FSC}$ model (0/0.143/0.5) | 2.8/3.1/3.6 | 2.9/3.2/3.7 |
| **Map-to-model CC** | | |
| Mask | 0.78 | |
| Box | 0.88 | |
| Peaks | 0.79 | |
| Volume | 0.79 | |
| **Composition** | | |
| Atoms | 6276 | |
| Amino acids | 787 | |
| Other ligands | 0 | |
| ***B*-factors** | | |
| Protein (min/max/mean) | 26.4/122.1/63.4 | |
| **r.m.s.d. from ideal values** | | |
| Bond length (Å) | 0.01 | |
| Bond angle (°) | 0.84 | |
| **Validation** | | |
| MolProbity score (Williams et al, 2018) | 2.89 | |
| Clashscore | 21.55 | |
| **Ramachandran plot** | | |
| Favored (%) | 85.0 | |
| Allowed (%) | 14.0 | |
| Outliers (%) | 1.0[a] | |
| **Rotamer outliers (%)** | 3.4 | |
| **CaBLAM outliers (%)** | 8.9 | |

[a]Ramachandran outliers (estimated by Phenix (Liebschner et al, 2019) cryo-EM validation tool and RAMPAGE (Lovell et al, 2003)): 8. The use of a different tool, employing different thresholds, could yield a different estimate.

| **Table S3 – Poorly-defined areas in the cryo-EM map** | | |
|---|---|---|
| **Residue range** | **Domain** | **Description** |
| 15-19 | BTF-1 | Subdomain α, loop forming the hypothetical sugar-binding site $\alpha_1$ |
| 24-29 | BTF-1 | Subdomain α, loop supporting the hypothetical sugar-binding site $\gamma_1$ |
| 355-360 | BTF-3 | Subdomain β, surface loop at the interface with domain BTF- 1 |
| 458-466 | BTF-4 | Subdomain β, surface loop |
| 522-529 | BTF-4 | Subdomain γ, surface loop |
| 531-548 | BTF-4 | Subdomain γ, loop forming the hypothetical sugar-binding site $\gamma_4$ |
| 578-589 | DUF | Loop connecting DUF strand-2 with DUF helix-1 |
| 603-612 | DUF | Loop connecting DUF helix-1 with DUF helix-2, supported by domain BTF-1 |
| 676-683 | DUF | Loop connecting DUF helix-4 (part of DUF insert #2) with DUF strand-4 |
| 756-770 | DUF | Loop connecting DUF strand-7 with helix-6 |

### Table S4 – CCTX paralogue domain boundaries and features outline

| Orthologue | | | | CCTX1 | CCTX2 | CCTX3 |
|---|---|---|---|---|---|---|
| **GenBank ID** | | | | CC1G_10075 | CC1G_10077 | CC1G_10083 |
| **UniProt ID** | | | | A8NDT5 | A8NDT7 | A8NDU3 |
| **Model source** | | | | Robetta homology model | cryo-EM | Robetta homology model |
| **Protein length (AA)** | | | | 774 | 787 | 783 |
| **BTF domain 1** | **Fold** | | | ricin B chain-like | ricin B chain-like | ricin B chain-like |
| | **Boundaries** | **Domain** | | 1-150 | 1-150 | 1-150 |
| | | **Subdomain α** | | 1-56 | 1-56 | 1-56 |
| | | **Subdomain β** | | 57-107 | 57-107 * | 57-107 |
| | | **Subdomain γ** | | 108-150 | 108-150 * | 108-150 |
| | **(QxW)₃** | **Subdomain α** | **Q** | Q 48 | Q 48 | Q 48 |
| | | | **W** | W 50 | W 50 | W 50 |
| | | **Subdomain β** | **Q** | Q 96 | Q 96 | Q 96 |
| | | | **W** | W 98 | W 98 | W 98 |
| | | **Subdomain γ** | **Q** | Q 45 | Q 145 | Q 145 |
| | | | **W** | W 147 | W 147 | W 147 |
| **Linker 1-2** | **Boundaries** | | | 151-156 | 151-156 | 151-156 |
| **BTF domain 2** | **Fold** | | | ricin B chain-like | ricin B chain-like | ricin B chain-like |
| | **Boundaries** | **Domain** | | 157-294 | 157-296 | 157-294 |
| | | **Subdomain α** | | 157-204 | 157-204 * | 157-204 |
| | | **Subdomain β** | | 205-252 | 205-254 | 205-252 |
| | | **Subdomain γ** | | 253-294 | 255-296 | 253-294 |
| | **(QxW)₃** | **Subdomain α** | **Q** | Q 196 | Q 196 | Q 196 |
| | | | **W** | W 198 | W 198 | W 198 |
| | | **Subdomain β** | **Q** | Q 244 | Q 244 | Q 244 |
| | | | **W** | W 246 | W 246 | W 246 |
| | | **Subdomain γ** | **Q** | Q 289 | Q 291 | Q 289 |
| | | | **W** | W 291 | W 293 | W 291 |
| **Linker 2-3** | **Boundaries** | | | 295-298 | 297-300 | 295-298 |
| **BTF domain 3** | **Fold** | | | ricin B chain-like | ricin B chain-like | ricin B chain-like |
| | **Boundaries** | **Domain** | | 299-435 | 301-436 | 299-436 |
| | | **Subdomain α** | | 299-345 | 301-347 | 299-346 |
| | | **Subdomain β** | | 346-393 | 348-396 | 347-394 |
| | | **Subdomain γ** | | 394-435 | 397-436 | 395-436 |
| | **(QxW)₃** | **Subdomain α** | **Q** | V 337 | V 339 | V 337 |
| | | | **W** | W 339 | W 341 | W 339 |
| | | **Subdomain β** | **Q** | D 383 | D 385 | D 383 |
| | | | **W** | W 385 | W 387 | R 385 |
| | | **Subdomain γ** | **Q** | R 430 | R 431 | R 430 |
| | | | **W** | W 432 | W 433 | W 432 |
| **Linker 3-4** | **Boundaries** | | | 436-442 | 437-443 | 437-442 |
| **BTF 4 domain** | **Fold** | | | ricin B chain-like | ricin B chain-like | ricin B chain-like |
| | **Boundaries** | **Domain** | | 443-552 | 444-554 | 443-552 |
| | | **Subdomain α** | | 443-484 | 443-487 | 443-485 |
| | | **Subdomain β** | | 485-513 | 488-516 | 486-514 |
| | | **Subdomain γ** | | 514-552 | 517-555 | 515-552 |
| | **(QxW)₃** | **Subdomain α** | **Q** | A 475 | A 478 | A 476 |
| | | | **W** | W 477 | W 480 | W 478 |
| | | **Subdomain β** | **Q** | F 506 | A 507 | A 505 |
| | | | **W** | L 508 | W 509 | W 507 |
| | | **Subdomain γ** | **Q** | Q 547 | Q 549 | Q 547 |
| | | | **W** | W 549 | W 551 | F 549 |
| **Linker 4-5** | **Boundaries** | | | 553-554 | 555-556 | 553-554 |
| **DUF domain 5** | **Fold** | | | Unknown (α+β) | Unknown (α+β) | Unknown (α+β) |
| | **Boundaries** | | | 555-774 | 557-787 | 555-783 |
| | **Kexin cleavage site** | | | 586-587 | 596-597 | 591-592 |

**Table S5 - CCTX2 β-trefoil fold (BTF) domains**
**Sequence identity (%) and structural similarity (r.m.s.d., Å, Cα alignment)**

| vs. | **BTF domain 1** | **BTF domain 2** | **BTF domain 3** | **BTF domain 4** |
|---|---|---|---|---|
| **BTF domain 1** | - | 35.8% / 1.9 Å | 9.8% / 2.5 Å | 14.2% /2.9 Å |
| **BTF domain 2** | 35.8% / 1.9 Å | - | 11.7% /2.7 Å | 17.1% /2.8 Å |
| **BTF domain 3** | 9.8% / 2.5 Å | 11.7% /2.7 Å | - | 18.4% /3.4 Å |
| **BTF domain 4** | 14.2% /2.9 Å | 17.1% /2.8 Å | 18.4% /3.4 Å | - |

### Table S6 – BTF domain homology

#### BTF domain 1

| PDB ID | chain | Z-score | r.m.s.d. | aligned residues | total protein length | % identity | Protein name | Organism | Function | UniProt | Reference |
|---|---|---|---|---|---|---|---|---|---|---|---|
| 3PG0 | A | 23.8 | 1.3 | 135 | 140 | 26 | synthetic β-threefoil protein | synthetic | proof-of-concept | synthetic | (Broom et al, 2012) |
| 8BAD | B | 23.1 | 1.5 | 140 | 351 | 15 | Tpp80Aa1 toxin | *Bacillus thuringiensis* | chimerolectin/ toxin | A0A4V1G8Q1 | (Best et al, 2022) |
| 3EF2 | A | 22.8 | 1.9 | 143 | 292 | 27 | *Marasiums oreades* agglutinin (MOA) | *Marasmius oreades* | chimerolectin/proteolytic toxin | Q8X123 | (Grahn et al, 2007) |
| 3VSF | C | 22.8 | 1.3 | 138 | 482 | 25 | 1,3Gal43A | *Acetivibrio thermocellus ATCC 27405* | exo-β-1,3-galactanase | A3DD67 | (Jiang et al, 2012) |
| 2X2T | A | 21.7 | 1.8 | 140 | 152 | 17 | *Sclerotinia sclerotiorum* agglutinin (SSA) | *Sclerotinia sclerotiorum* | lectin | A7XUK7 | (Sulzenbacher et al, 2010) |
| 3NBE | B | 21.4 | 1.9 | 138 | 147 | 25 | *Clitocybe nebularis* ricin B-like lectin (CNL) | *Clitocybe nebularis* | lectin | B2ZRS9 | (Pohleven et al, 2012) |
| 3PHZ | B | 21.3 | 1.7 | 135 | 278 | 27 | *Polyporus squamosus* lectin 1a (PSL1a) | *Polyporus squamosus* | chimerolectin/proteolytic toxin | Q75WT9 | (Kadirvelraj et al, 2011) |
| 4G9M | A | 20.9 | 1.6 | 132 | 142 | 20 | *Rhizoctonia solani* agglutinin | *Rhizoctonia solani* | lectin | L8WGI4 | (Skamnaki et al, 2013) |
| 2VSA | A | 20.3 | 1.7 | 135 | 822 | 21 | Mosquitocidal holotoxin (MTX) | *Bacillus sphaericus* | chimerolectin/toxin | Q03988 | (Treiber et al, 2008) |
| 1V6W | A | 20.1 | 1.7 | 128 | 436 | 24 | xylanase | *Streptomyces olivaceoviridis* | endo-1,4-β-D-xylanase | Q7SI98 | (Fujimoto et al, 2004) |

#### BTF domain 2

| PDB ID | chain | Z-score | r.m.s.d. | aligned residues | total protein length | % identity | Protein name | Organism | Function | UniProt | Reference |
|---|---|---|---|---|---|---|---|---|---|---|---|
| 3PG0 | A | 23.0 | 1.8 | 134 | 140 | 26 | synthetic β-threefoil protein | synthetic | proof-of-concept | synthetic | (Broom et al, 2012) |
| 3VSF | E | 22.6 | 1.7 | 134 | 461 | 28 | 1,3Gal43A | *Acetivibrio thermocellus ATCC 27405* | exo-β-1,3-galactanase | A3DD67 | (Jiang et al, 2012) |
| 2IHO | A | 22.0 | 2.2 | 139 | 292 | 27 | *Marasiums oreades* agglutinin (MOA) | *Marasmius oreades* | chimerolectin/proteolytic toxin | Q8X123 | (Grahn et al, 2007) |
| 8BAD | B | 21.9 | 1.7 | 125 | 351 | 19 | Tpp80Aa1 toxin | *Bacillus thuringiensis* | chimerolectin/ toxin | A0A4V1G8Q1 | (Best et al, 2022) |
| 3A21 | A | 20.7 | 1.6 | 125 | 614 | 29 | β-L-arabinopyranosidase | Streptomyces avermitilis | β-L-arabinopyranosidase | Q82L26 | (Ichinose et al, 2009) |
| 4G9M | A | 20.3 | 1.8 | 128 | 142 | 27 | *Rhizoctonia solani* agglutinin | *Rhizoctonia solani* | lectin | L8WGI4 | (Skamnaki et al, 2013) |
| 6IFA | C | 20.2 | 1.6 | 124 | 126 | 22 | β-trefoil lectin (EntTref) | *Entamoeba histolytica* HM-1:IMSS | lectin | N9TFI9 | (Khan et al, 2020) |
| 5GQD | A | 20.2 | 1.6 | 122 | 427 | 27 | Glycoside Hydrolase Family 10 Xylanase | *Streptomyces olivaceoviridis* | xylanase | Q7SI98 | (Suzuki et al, 2018) |
| 3NBE | B | 19.7 | 2.5 | 133 | 147 | 18 | *Clitocybe nebularis* ricin B-like lectin (CNL) | *Clitocybe nebularis* | lectin | B2ZRS9 | (Pohleven et al, 2012) |
| 2VLC | B | 19.6 | 1.7 | 123 | 518 | 18 | Cinnamomum camphora | *Cinnamomum camphora* | ribosome inactivating protein | Q94BW3 | (Azzi et al, 2009) |

#### BTF domain 3

| PDB ID | chain | Z-score | r.m.s.d. | aligned residues | total protein length | % identity | Protein name | Organism | Function | UniProt | Reference |
|---|---|---|---|---|---|---|---|---|---|---|---|
| 3AJ6 | A | 19.5 | 1.9 | 130 | 283 | 19 | botulinum type C progenitor toxin | *Clostridium botulinum* | toxin | P0DPR0 | (Nakamura et al, 2011) |
| 2X2S | B | 19.1 | 2.0 | 133 | 152 | 12 | *Sclerotinia sclerotiorum* agglutinin (SSA) | *Sclerotinia sclerotiorum* | lectin | A7XUK7 | (Sulzenbacher et al, 2010) |
| 3EF2 | A | 19.1 | 2.1 | 134 | 292 | 11 | *Marasiums oreades* agglutinin (MOA) | *Marasmius oreades* | chimerolectin/proteolytic toxin | Q8X123 | (Grahn et al, 2009) |
| 1QXM | A | 19.1 | 1.9 | 130 | 283 | 19 | hemagglutinin component (HA1) | *Clostridium botulinum* D phage | lectin | P0DPR1 | (Inoue et al, 2003) |
| 3O49 | A | 18.8 | 1.7 | 122 | 123 | 17 | Symfoil-1 | synthetic | proof-of-concept | synthetic | (Lee & Blaber, 2011) |
| 3VSF | E | 18.6 | 1.9 | 126 | 461 | 17 | 1,3Gal43A | Acetivibrio thermocellus ATCC 27405 | exo-β1,3-galactanase | A3DD67 | (Jiang et al, 2012) |
| 1UPS | B | 18.2 | 2.1 | 125 | 397 | 21 | GlcNAcα1-4Gal | Clostridium perfringens | endo-β-galactosidase | Q934G8 | (Tempel et al, 2005) |
| 3PHZ | B | 18.1 | 1.9 | 127 | 278 | 17 | *Polyporus squamosus* lectin 1a (PSL1a) | *Polyporus squamosus* | chimerolectin/proteolytic toxin | Q75WT9 | (Nakamura et al, 2008) |
| 7KCG | A | 18.1 | 1.7 | 124 | 143 | 19 | mosquito salivary protein | *Culex quinquefasciatus* | lectin (probable) | Q6TRZ5 | (Kern et al, 2021) |
| 2VSA | A | 18.0 | 2.0 | 127 | 822 | 13 | Mosquitocidal holotoxin (MTX) | *Bacillus sphaericus* | chimerolectin/toxin | Q03988 | (Treiber et al, 2008) |

#### BTF domain 4

| PDB ID | chain | Z-score | r.m.s.d. | aligned residues | total protein length | % identity | Protein name | Organism | Function | UniProt | Reference |
|---|---|---|---|---|---|---|---|---|---|---|---|
| 3O4D | A | 16.2 | 1.6 | 104 | 123 | 16 | Symfoil-4P | synthetic | proof-of-concept | synthetic | (Lee & Blaber, 2011) |
| 4G9N | A | 15.4 | 1.8 | 109 | 142 | 21 | Rhizoctonia solani agglutinin | *Rhizoctonia solani* | lectin | L8WGI4 | (Skamnaki et al, 2013) |
| 3VSF | A | 15.4 | 1.7 | 108 | 461 | 17 | 1,3Gal43A | *Acetivibrio thermocellus ATCC 27405* | exo-β-1,3-galactanase | A3DD67 | (Jiang et al, 2012) |
| 1BFC | A | 15.0 | 2.0 | 106 | 124 | 17 | bFGF - basic fibroblast growth factor/FGF2 | *Homo sapiens* | growth factor | P09038 | (Faham et al, 1996) |
| 7ZNX | A | 14.9 | 2.1 | 107 | 136 | 20 | cocaprin 1 | *Coprinopsis cinerea* | cysteine and aspartic proteases inhibitor | A8PCJ3 | (Renko et al, 2022) |
| 3UD7 | C | 14.9 | 2.1 | 107 | 127 | 13 | aFGF - basic fibroblast growth factor/FGF1 | *Homo sapiens* | hormone | P05230 | (Hu et al, 2012) |
| 3AJ6 | A | 14.7 | 1.7 | 106 | 283 | 14 | HA33/C (HA1) module of type C neurotoxin | *Clostridium botulinum* | toxin | P0DPR0 | (Nakamura et al, 2011) |
| 4IYB | B | 14.5 | 1.9 | 107 | 140 | 15 | Macrolepiota procera ricin B-like lectin (MPL) | *Macrolepiota procera* | lectin | F6KMV5 | (Žurga et al, 2014) |
| 8BAD | A | 14.5 | 1.8 | 109 | 350 | 11 | Tpp80Aa1 | *Bacillus thuringiensis* | toxin | A0A4V1G8Q1 | (Best et al, 2022) |
| 2VSE | B | 14.5 | 1.7 | 106 | 822 | 16 | Mosquitocidal holotoxin (MTX) | *Bacillus sphaericus* | chimerolectin/toxin | Q03988 | (Treiber et al, 2008) |

**Table S7. Plasmids generated in this study**

| Construct | Usage | Toxicity | Reference |
|---|---|---|---|
| pET24_CCTX2 | Purification of CCTX2 (functional studies) | toxic | This study |
| pET24_CCTX1 | Purification of CCTX1 | toxic | This study |
| pET24_CCTX3 | Purification of CCTX3 | toxic | This study |
| pET24_CCTX2-ΔN | Purification of CCTX2ΔN deletion mutant (lacking residues 2-303) | non-toxic | This study |
| pET24_CCTX2-ΔC | Purification of CCTX2ΔC deletion mutant (lacking residues 570-787) | non-toxic | This study |
| pET24_CCTX2-ΔHSEL | Purification of CCTX2ΔHSEL deletion mutant (lacking residues 784-787) | non-toxic | This study |
| pET22_MOA | Purification of MOA | toxic | (Wohlschlager et al, 2011) |
| pET24_CCL2 | Purification of CCL2 | toxic | (Schubert et al, 2012) |
| pET24_CGL2 | Purification of CGL2 | toxic | (Butschi et al, 2010) |
| pET22b(+)-His8-TEVcs-CCTX2 | Purification of CCTX2 (structural studies) | not tested | This study |
| pET24_CCTX2-D567A | Purification of CCTX2 D567A | toxic | This study |
| pET24_CCTX2-D572A | Purification of CCTX2 D572A | toxic | This study |
| pET24_CCTX2-D585A | Purification of CCTX2 D585A | non-toxic up to 500 µg/ml | This study |
| pET24_CCTX2-K600A | Purification of CCTX2 K600A | toxic | This study |
| pET24_CCTX2-Q635A | Purification of CCTX2 Q635A | non-toxic up to 100 µg/ml | This study |
| pET24_CCTX2-D639A | Purification of CCTX2 D639A | toxic | This study |
| pET24_CCTX2-E656A | Purification of CCTX2 E656A | non-toxic up to 100 µg/ml | This study |
| pET24_CCTX2-E658A | Purification of CCTX2 E658A | toxic | This study |
| pET24_CCTX2-R660A | Purification of CCTX2 R660A | non-toxic up to 50 µg/ml | This study |
| pET24_CCTX2-R685A | Purification of CCTX2 R685A | non-toxic up to 50 µg/ml | This study |
| pET24_CCTX2-D687A | Purification of CCTX2 D687A | non-toxic | This study |
| pET24_CCTX2-Q689A | Purification of CCTX2 Q689A | non-toxic | This study |
| pET24_CCTX2-R724A | Purification of CCTX2 R724A | non-toxic up to 200 µg/ml | This study |
| pET24_CCTX2-R732A | Purification of CCTX2 R732A | toxic | This study |
| pET24_CCTX2-T739A | Purification of CCTX2 T739A | non-toxic up to 500 µg/ml | This study |
| pET24_CCTX2-N767A | Purification of CCTX2 N767A | toxic | This study |
| pET24_CCTX2-DxQ | Purification of CCTX2 D687A, Q689A double mutant | non-toxic | This study |
| pET24_CCTX2-RxDxQ | Purification of CCTX2 R685A, D687A, Q689A triple mutant | non-toxic | This study |
| pET24_CCTX2-DDDxL | Purification of CCTX2 D572A, D573A, D574A, L576A quadruple mutant | non-toxic up to 100 µg/ml | This study |
| pET24_CCTX2-KR | Purification of CCTX2 E595A, K596A, R597A triple mutant | non-toxic up to 100 µg/ml | This study |

**Table S8.** ***Caenorhabditis*** **and** ***Escherichia coli*** **strains used in this study.**

| Strain name | Genotype | Feature | Reference |
|---|---|---|---|
| ***C. elegans*** | | | |
| VC424(BEC-1) | *bec-1(ok700) IV/nT1 [qIs51]* | *bec-1* (VPS-34-recruiting protein) deletion mutant balanced by GFP-marked translocation | CGC |
| HY494(BRE-2) | *bre-2(ye31)* | *bre-2* (β1,3 galactosyltransferase) deletion mutant | CGC |
| HY483(BRE-3) | *bre-3(ye26)* | *bre-3* (β1,4 mannosyltransferase) deletion mutant | CGC |
| HY498(BRE-5) | *bre-5(ye17)* | *bre-5* (β1,3 *N*-acetylglucosamine transferase) deletion mutant | CGC |
| HY485(BRE-4) | *bre-4(ye27)* | *bre-4* (β1, 4 *N*-acetylgalactosamine transferase) deletion mutant | CGC |
| GK70 | *dkIs37[$P_{act\text{-}5}$::GFP:pgp-1]* | PGP-1 fused to GFP | CGC |
| N2 (Bristol type) | | Wild type | CGC |
| RT1315 | *pwIs503[vha- 6p::mans::GFP +Cbr-unc-119(+)* | MANS (α-Mannosidase II) fused to GFP | CGC |
| RT311 | *pwIs69[vha6p::GFP::rab-11 + unc-119(+)]* | RAB-11 fused to GFP | CGC |
| RT327 | *pwIs72[vha-6p::GFP::rab-5 + Cbr-unc-199(+)]* | RAB-5 fused to GFP | CGC |
| VC625(VPS-52) | *vps-52(ok853)* | *vps-52* (GARP complex) deletion mutant | CGC |
| VC985(VPS-54) | *vps-54(ok1463) V* | *vps-54* (GARP complex) deletion mutant | CGC |
| ***C. briggsae*** | | | |
| AF16 | | | CGC |
| ***C. tropicalis*** | | | |
| JU1373 | | | CGC |
| ***E. coli*** | | | |
| DH5α | $F^-$ *ϕ80lacZΔM15 Δ(lacZYA-argF)U169 recA1 endA1 hsdR17(rK$^-$, mK+) phoA supE44 λ$^-$ thi-1 gyrA96 relA1* | Cloning | Thermo Fisher Scientific |
| Turbo | F' *proA$^+$B$^+$ lacI$^q$* Δ *lacZ* M15/ *fhuA2* *Δ(lac-proAB) glnV gal R(zgb- 210*::*Tn10*)$Tet^S$ *endA1 thi-1 Δ(hsdS-mcrB)5* | Cloning | New England Biolabs |
| OP50 | *ura-* | *C. elegans* feeding | CGC |
| BL21(DE3) | *$F^-$ ompT hsdSb($r_B^-$$m_B^-$) gal dcm* (DE3) F` (traD36, *pro*AB+ *lac*Iq, Δ(*lac*Z)M15) | Protein production | New England Biolabs |
| C41(DE3) | $F^-$ *ompT hsd*$S_B$ ($r_B^-$ $m_B^-$) *gal dcm* Δ *AcrB* (DE3) | Protein production | Sigma-Aldrich Merck |

**Note**: CGC: Caenorhabditis Genomics Center (CGC), University of Minnesota, Minneapolis USA

**Table S9. Selected primers used in this study**

| Primer | Sequence | Usage |
|---|---|---|
| CCTX2for_NdeI<br>CCTX2rev_NotI | GGAGTCGGCATATGGCTCTCAACGAAGGTG<br>GAATAGCGGCCGCCTACAACTCGGAGTGCTTG | Cloning of CCTX2 cDNA |
| CCTX1for_His_NdeI<br><br>CCTX1rev_NotI | GCCCGCCATATGCATCATCATCATCATCATTCTATCTCCGAA GGCGTTTACTGG<br>GTAATAGCGGCCGCCTACAACTCGGAGTG | Cloning of CCTX1 cDNA |
| CCTX3for_His_NdeI<br><br>CCTX3rev_NotI | GCCCGGCCATATGCATCATCATCATCATCATTCTCTTTCTCA GGGTGTTTACTGG<br>ATATAGCGGCCGCCTAGAGCTCGGAGTGCTTG | Cloning of CCTX3 cDNA |
| CCTX2ΔNfor_His_NdeI<br><br>CCTX2ΔC_rev_NotI<br><br>CCTX2ΔHSEL_rev_NotI | CTGGGTACATATGCATCATCATCATCATCATCATCATCCCCT GCCCTCTCCTGGTCC<br>GTGATGCGGCCGCCTAATAGATCAGATCGTAGTCATTGC<br><br>GAATAGCGGCCGCCTACTTGATTGCCGAAGTACCGACG | Cloning of CCTX2ΔN (truncation of the two N-terminal BTF domains)<br>Cloning CCTX2ΔC (truncation of the C-terminal DUF domain)<br>Cloning of CTX2ΔHSEL (truncation of the C-terminal HSEL sequon, CCTX2for_NdeI was used as for primer) |
| CCTX2_fwd<br>CCTX2_rev<br>pET22_fwd<br>pET22_rev | CTTCCAATCCATGGCTCTCAACGAAGGTG<br>CGATATCCATCTACAACTCGGAGTGCTTG<br>CGAGTTGTAGATGGATATCGGAATTAATTCG<br>TGAGAGCCATGGATTGGAAGTACAAGTTTTC | Cloning of full-length CCTX2 including a N-terminal $His_8$ tag followed by a TEV cleavage site into pET22b |
